# Periphilin self-association underpins epigenetic silencing by the HUSH complex

**DOI:** 10.1101/2019.12.18.881300

**Authors:** Daniil M. Prigozhin, Anna Albecka, Christopher H. Douse, Iva A. Tchasovnikarova, Richard T. Timms, Laura E. Farleigh, Shun Oda, Stefan M. V. Freund, Sarah Maslen, Paul J. Lehner, Yorgo Modis

## Abstract

Transcription of integrated DNA from viruses or transposable elements is tightly regulated to prevent pathogenesis. The Human Silencing Hub (HUSH), composed of Periphilin, TASOR and MPP8, silences transcriptionally active viral and endogenous transgenes. HUSH recruits effectors that alter the epigenetic landscape and chromatin structure, but how HUSH recognizes target loci and represses their expression remains unclear. We identify the physicochemical properties of Periphilin necessary for HUSH assembly and silencing. A disordered N-terminal domain (NTD) and structured C-terminal domain are essential for silencing. A crystal structure of the Periphilin-TASOR core complex shows Periphilin forms *α*-helical homodimers, which each bind a single TASOR molecule. The NTD binds RNA non-specifically and forms insoluble aggregates through an arginine/tyrosine-rich sequence reminiscent of low-complexity regions from self-associating RNA-binding proteins. Residues required for TASOR binding and aggregation were required for HUSH-dependent silencing and genome-wide deposition of repressive mark H3K9me3. The NTD was functionally complemented by low-complexity regions from certain RNA-binding proteins and proteins that form condensates or fibrils. Our work suggests the associative properties of Periphilin promote HUSH aggregation on nascent transcripts.

## Introduction

More than half of the human genome consists of transposable elements (TEs). TEs have evolved to fulfill important cellular functions. TEs drive the evolution of transcriptional networks by spreading transcription factor binding sites, promoters and other regulatory elements (Chuong *et al*, 2016; Friedli & Trono, 2015). TE-derived regulatory elements are particularly important in embryogenesis, when global hypomethylation promotes transcription. Key pluripotency-associated transcription factors involved in cell fate determination bind to sites within TEs (Friedli & Trono, 2015). TE genes also serve as a genetic reservoir that can be coopted by the host. For example, TE-derived proteins catalyze V(D)J recombination (Zhou *et al*, 2004) and syncytiotrophoblast fusion in placental development (Dupressoir *et al*, 2012; Friedli & Trono, 2015).

A subset of TEs can autonomously replicate through an RNA intermediate and reintegrate into the genome like retroviruses. Some of these TEs are endogenous retrovirus (ERV) genomes inherited from ancestral infections of germline. The other type of autonomously replicating TE in humans are the non-viral LINE-1 (long interspersed nuclear element-1) retroelements. Active ERVs and LINE-1s are transcribed and encode reverse transcriptase and integrase enzymes, which convert the transcripts into DNA and reintegrate it into the host genome (Friedli & Trono, 2015). This amplifying retrotransposition mechanism has allowed ERVs and L1s to accumulate in the human genome. Approximately 100 human LINEs are replication-competent and cause new integration events in 2-5% of the population (Friedli *et al*, 2014; Goodier, 2016).

Transcription and retrotransposition of TEs must be tightly regulated to prevent harmful gene expression and genome damage. Accumulation of TE transcripts is associated with autoimmune diseases including geographic atrophy, lupus and Sjögren’s syndrome (Goodier, 2016; Hung *et al*, 2015). Aberrant expression of proteins from the human ERV HERV-K is associated with cancer and neurodegeneration (Li *et al*, 2015). Reactivation of ERVs and LINE-1s in somatic cells is also associated with cancer, through disruption of tumor suppressor genes or enhanced transcription of oncogenes (Hancks & Kazazian, 2016; Lamprecht *et al*, 2010). Disruption of protein coding sequences by transposition events is additionally linked to genetic disorders such as hemophilia and cystic fibrosis (Hancks & Kazazian, 2016; Lamprecht *et al*, 2010).

A central mechanism cells have evolved to control potentially pathogenic expression and transposition of TEs and infectious viruses alike is epigenetic silencing. Among the most important sources of epigenetic silencing in humans is the Human Silencing Hub (HUSH) complex, consisting of three proteins: Periphilin, TASOR and MPP8 (Tchasovnikarova *et al*, 2015). HUSH silences the genomes from newly integrated lentiviruses (Tchasovnikarova *et al*, 2015), and from unintegrated retroviruses via the DNA-binding protein NP220 (Zhu *et al*, 2018). Vpr and Vpx proteins from lentiviruses including HIV target HUSH for proteasomal degradation, demonstrating the importance of HUSH-dependent silencing in controlling lentiviral infection (Chougui *et al*, 2018; Greenwood *et al*, 2019; Yurkovetskiy *et al*, 2018). HUSH also silences hundreds of transcriptionally active or recently integrated genes, with a degree of selectivity for full-length LINE-1s located in euchromatic environments, often within introns of actively transcribed genes (Liu *et al*, 2018). How HUSH recognizes integrated target loci and represses their expression remains unclear. A necessary component of HUSH-dependent silencing is deposition of histone H3 lysine 9 trimethylation (H3K9me3), a transcriptionally repressive mark, through recruitment of the H3K9 methyltransferase SETDB1 and its stabilizing factor ATF7IP (Tchasovnikarova *et al*, 2015; Timms *et al*, 2016). HUSH silencing also requires MORC2, a DNA-binding ATPase that acts as a chromatin remodeler (Douse *et al*, 2018; Tchasovnikarova *et al*, 2017).

All three HUSH subunits are essential for silencing (Tchasovnikarova *et al*, 2015) but their biochemical and structural properties remain unknown. Hence, how the HUSH complex assembles, recognizes its target loci and modifies chromatin to repress gene expression remains unclear. In this study, we delineate the key structural and physicochemical attributes of Periphilin and how they contribute to HUSH function. Periphilin was originally identified as a highly insoluble nuclear protein cleaved by caspase-5 (Kazerounian & Aho, 2003). N-terminal sequences contain the determinants for insolubility and the C-terminal region contains predicted *α*-helical heptad repeats proposed to form dimers based on a yeast two-hybrid assay (Kazerounian & Aho, 2003). Periphilin is indispensable for development. In mice, homozygous deficiency of Periphilin is lethal early in embryogenesis and heterozygous deficiency is compensated by increased expression from the wild-type allele (Soehn *et al*, 2009). Overexpression of Periphilin transcriptionally represses certain proteins causing cell cycle arrest (Kurita *et al*, 2007; Kurita *et al*, 2004). Isoform 2 of Periphilin, one of at least 8 isoforms, was identified in a gene-trap mutagenesis screen as a component of the HUSH complex that binds TASOR but not MPP8 (Douse *et al*, 2019; Tchasovnikarova *et al*, 2015). Curiously, some of the isoform diversity is driven by TE insertion into Periphilin coding sequences (Huh *et al*, 2006). Periphilin was also identified as an mRNA-binding protein in a screen of the protein-mRNA interactome in proliferating human cells (Castello *et al*, 2012). Here, we report the crystal structure of a minimal Periphilin-TASOR complex and identify the key physicochemical properties of Periphilin necessary for HUSH complex assembly and epigenetic silencing. The Periphilin C-terminal region directs HUSH complex assembly by dimerizing and binding a single TASOR molecule through *α*-helical coiled-coil interactions. A disordered N-terminal domain (NTD) binds RNA non-specifically and mediates self-aggregation through a sequence enriched in arginine and tyrosine residues. The sequence of the NTD is reminiscent of— and functionally complemented by— low-complexity regions from RNA-binding proteins, and from certain proteins that self-associate to form biomolecular condensates or phase separations. Our findings suggest Periphilin may contribute to the recognition and co- or post-transcriptional repression of HUSH target loci by binding and sequestering nascent transcripts. This work provides a foundation to design strategies to control HUSH activity, with important potential therapeutic applications.

## Results

### Both N- and C-terminal regions of Periphilin are required for HUSH function

To identify the subdomains of Periphilin required for HUSH function, we generated various Periphilin constructs with N- or C-terminal truncations and assessed their silencing activity as part of the HUSH complex (**Fig. 1A**). We used the 374-amino acid isoform 2 of Periphilin (UniProt Q8NEY8-2) as the reference sequence in this study rather than the longer isoform 1 (UniProt Q8NEY8-1), as isoform 2 fully restores HUSH function in Periphilin-deficient cells (Tchasovnikarova *et al*, 2015). Repression of a lentiviral GFP reporter transduced into Periphilin knockout (Periphilin KO) HeLa cells was used as a measure of silencing activity. GFP fluorescence was detected by flow cytometry. As reported previously (Tchasovnikarova *et al*, 2015), GFP reporter expression was repressed in wild-type cells and derepressed in Periphilin KO cells (**Fig. 1B**). Transduction of Periphilin KO cells with a Periphilin construct lacking amino acids 1-24 (1Δ25) or 1-69 (1Δ70) rescued reporter expression to the same extent as transduction with wild-type Periphilin. However, transduction with Periphilin constructs lacking amino acids 351-374 (350Δ374) or 1-126 (1Δ127) failed to rescue reporter repression in Periphilin KO cells, indicating that both N- and C-terminal regions of Periphilin are required for HUSH function.

**Fig. 1.**
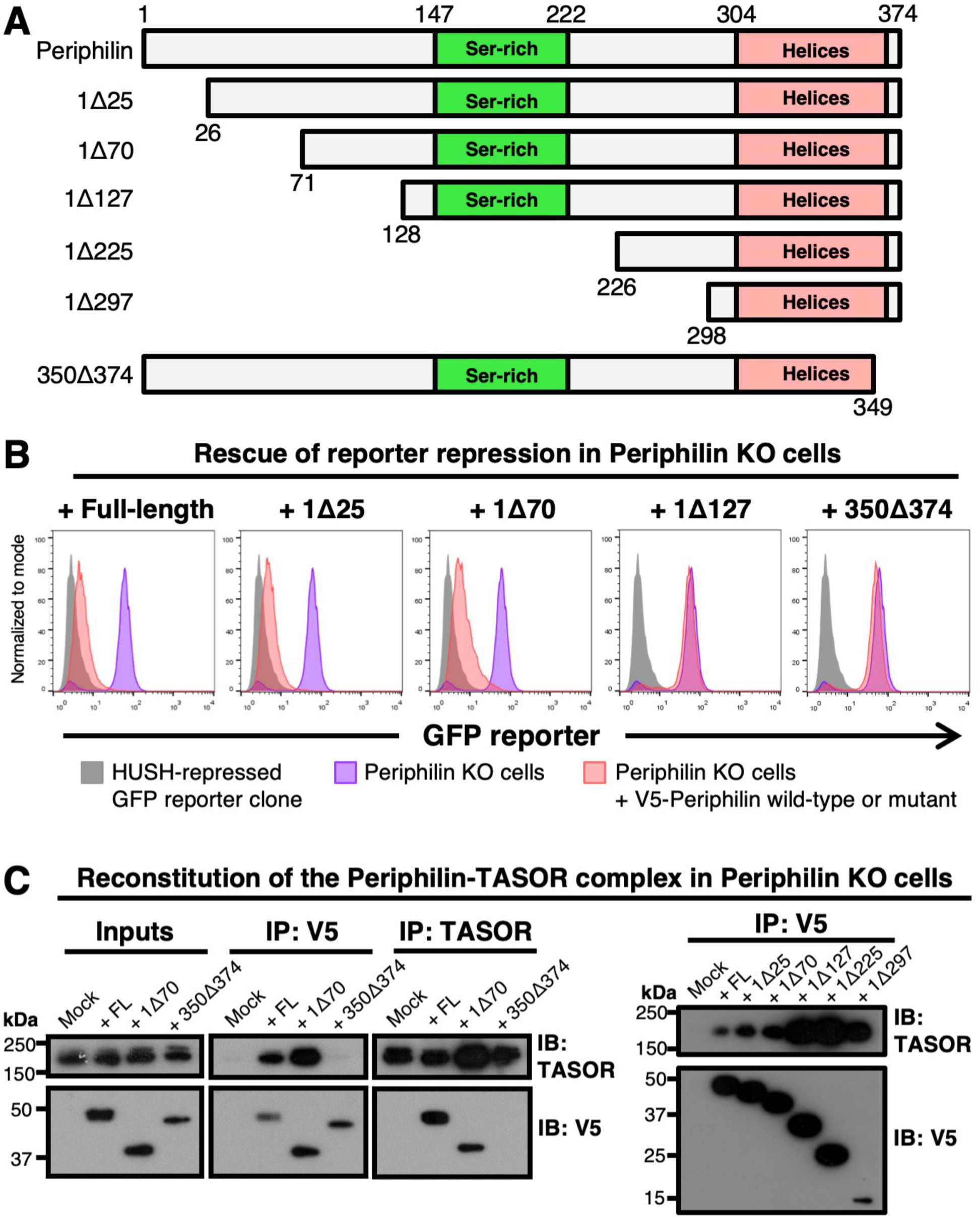
Both N- and C-terminal regions of Periphilin are required for HUSH function but only the C-terminal region is required for HUSH complex formation. (**A**) Schematic representation of Periphilin constructs used for complementation assay. All variants were expressed with an N-terminal V5 tag. (**B**) Repression of a lentiviral GFP reporter in Periphilin KO cells complemented with full-length Periphilin or truncation mutants (data shown 7 days post-transduction). The 1Δ127 and 350Δ374 variants fail to rescue reporter repression. (**C**) Pulldown assays with Periphilin and TASOR, the largest HUSH component. Periphilin and TASOR were immunoprecipitated (IP) from Periphilin KO cells complemented with Periphilin deletion mutants on Protein A/G resin decorated with anti-V5 or anti-TASOR primary antibody, respectively. TASOR and Periphilin proteins bound to the resin were quantified by Western immunoblot (IB). Only the C-terminal Periphilin deletion (350Δ374) abolished TASOR binding. The V5 tag was used to detect Periphilin.

The loss of HUSH function with the 350Δ374 and 1Δ127 Periphilin mutants could be due to loss of an intrinsic activity of Periphilin or failure of Periphilin to be recruited to the HUSH complex. To distinguish between these, we measured coprecipitation of Periphilin and TASOR in pulldown assays. Periphilin and TASOR were purified on an immunoaffinity resin (immunoprecipitated) from lysates of Periphilin KO cells complemented with Periphilin deletion mutants. TASOR coeluted with all N-terminal deletion mutants tested, up to 1Δ297 (**Fig. 1C**). Conversely, wild-type and 1Δ70 Periphilin both coeluted with TASOR. In contrast, TASOR did not associate with immunoprecipitated 350Δ374 Periphilin and 350Δ374 Periphilin did not associate with TASOR. Hence, only the C-terminal region of Periphilin is required for binding to TASOR, and the N-terminal region of Periphilin must have other properties necessary for HUSH function.

### Structure of the core Periphilin-TASOR complex identifies interfaces required for HUSH function

Having established that the C-terminal region of Periphilin (residues 297-374) is required for binding to TASOR, we sought to identify the structural determinants of Periphilin-TASOR assembly. We recently mapped the Periphilin binding region in TASOR to within residues 1000-1085 (Douse *et al*, 2019). Initial attempts to crystallize Periphilin-TASOR complexes failed until we determined that residues 285-291 and 368-374 of Periphilin were disordered from NMR spectra with ^15^N- and ^13^C-labeled Periphilin (**Fig. EV1**). Thus, a crystal structure of Periphilin residues 292-367 bound to TASOR residues 1014-1095 was determined using the single anomalous dispersion (SAD) phasing method, with bromine as the anomalous scatterer (**Table 1**). The structure contains two Periphilin molecules and a single TASOR molecule (**Fig. 2A**). The Periphilin fragments form helical hairpins with a mixture of *α*-helix and 3_10_-helix secondary structure. The two Periphilin hairpins pack against each other via a 118 Å^2^ hydrophobic interface formed by the hydrophobic side chains of Leu326, Leu333 and Ile337. The resulting Periphilin homodimer has twofold symmetry. The TASOR molecule forms two *α*-helices that wrap around the outer surfaces of the Periphilin dimer. The TASOR helices add a third helix to each Periphilin helical hairpin to form two three-helix coiled-coils. Each TASOR helix forms leucine zipper-type hydrophobic contacts, which typify helical coiled-coils. Unusually, however, each Periphilin subunit binds to a different TASOR sequence (residues 1014-1052 and 1072-1093, respectively) with an identical binding surface (**Fig. 2A**). Notably residues 1055-1071 of TASOR, between the two Periphilin-binding segments, are disordered, but these 17 residues could easily span the 35-40 Å trajectory needed to connect residues 1054 and 1072 in the Periphilin-TASOR complex. Binding of TASOR to Periphilin buries a total of 428 Å^2^. The 2:1 stoichiometry of the Periphilin-TASOR core complex was confirmed in solution by size-exclusion chromatography coupled with multiangle light scattering (SEC-MALS; **Fig. 2B**) and non-denaturing mass spectrometry (**Fig. EV2**).

**Fig. 2.**
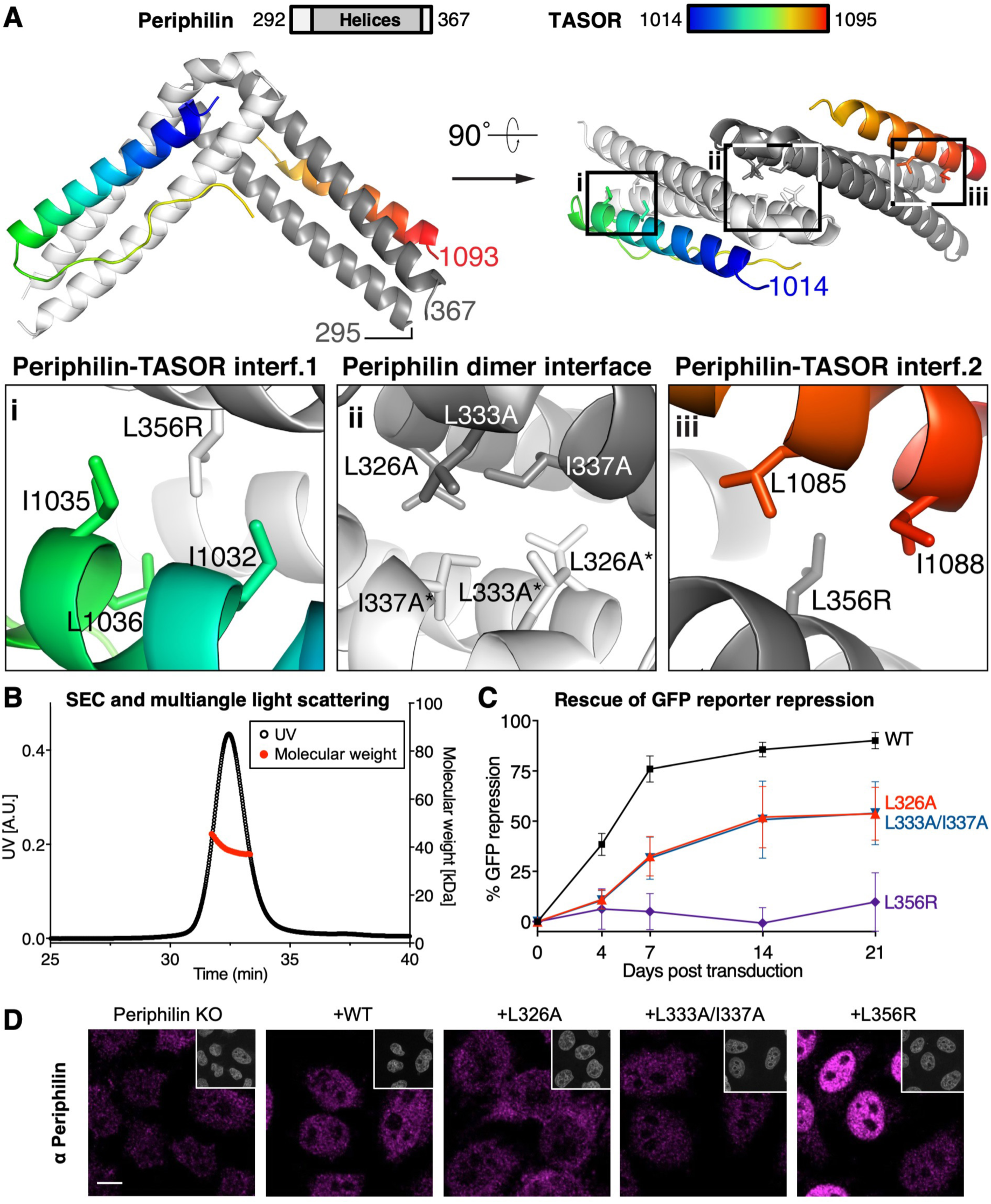
Periphilin and TASOR form a 2:1 complex required for HUSH function. (**A**) Crystal structure of the Periphilin-TASOR core complex. The Periphilin fragment (residues 292-367, light/dark grey) forms a homodimer of helical hairpins. The TASOR fragment (residues 1014-1095, rainbow colors) wraps around the Periphilin dimer, adding an *α*-helix to each Periphilin hairpin to form two helical coiled coils. Insets show close-up views of the Periphilin-TASOR interfaces (“i”, “iii”) and the Periphilin dimer interface (“ii”). Residues forming key contacts and mutations designed to disrupt Periphilin-TASOR complex formation are labeled. (**B**) SEC-MALS of Periphilin-TASOR core complex. The molecular weight calculated from light scattering data is consistent with a 2:1 complex in solution. (**C**) Repression of a lentiviral GFP reporter in Periphilin KO cells complemented with Periphilin mutants designed to inhibit Periphilin-TASOR complex assembly. Reporter expression was monitored over 21 days by flow cytometry. The log_10_(GFP fluorescence) data for live cells were converted to percent repression activity with wild-type HeLa set at 100% and Periphilin KO cells set a 0% repression (see Methods). (**D**) Immunofluorescence microscopy of Periphilin KO cells transduced with Periphilin mutants affecting Periphilin-TASOR complex assembly. Cells were fixed 4 days post-transduction and stained with anti-Periphilin antibody (magenta) and DAPI (grey, insets). Scale bar, 10 µm.

**Table 1.** Crystallographic data collection and refinement statistics for the Periphilin-TASOR complex.

| **Data collection** | **Native** | **Br derivative** |
| --- | --- | --- |
| X-ray source | DLS I04-1 | DLS I04-1 |
| Space group | *P*3<sub>2</sub>21 | *P*3<sub>2</sub>21 |
| Cell dimensions |  |  |
| a = b, c (Å) | 93.57, 84.97 | 93.96, 82.30 |
| α = β, γ (°) | 90, 120 | 90, 120 |
| Wavelength (Å) | 0.91587 | 0.91587 |
| Resolution (Å)<sup>a</sup> | 41.0-2.51 (2.58-2.51) | 81.4-2.93 (3.01-2.93) |
| Observations | 150,312 | 183,145 |
| Unique reflections | 15,065 | 9,350 |
| *R*<sub>merge</sub><sup>b</sup> | 0.081 (1.014) | 0.088 (2.426) |
| *R*<sub>pim</sub><sup>c</sup> | 0.029 (0.360) | 0.020 (0.576) |
| <\|*I*> / σ\| | 15.0 (2.1) | 25.5 (1.4) |
| Completeness (%) | 99.9 (100) | 100 (100) |
| Multiplicity | 5.1(4.9) | 10.5 (9.6) |
| CC(1/2) | 0.998 (0.735) | 0.999 (0.542) |
| **SAD Phasing with Br** |  |  |
| CC<sub>ano</sub> |  | 0.57 (0.02) |
| \|D<sub>ano</sub>\| / σD<sub>ano</sub> |  | 1.56 (0.71) |
| Figure of merit, FOM (initial) |  | 0.36 |
| FOM (density modified) |  | 0.43 |
| FOM (after autobuild) |  | 0.65 |
| **Refinement** |  |  |
| Resolution (Å) | 41.0-2.51 |  |
| *R*<sub>work</sub> / *R*<sub>free</sub><sup>d</sup> | 0.228 / 0.271 |  |
| No. of non-H atoms |  |  |
| Protein | 1689 |  |
| Sulfate ions | 1 |  |
| Water | 0 |  |
| No. riding H atoms | 1707 |  |
| Mean B-factor (Å<sup>2</sup>)<sup>e</sup> | 108 |  |
| MolProbity Clashscore | 3.23 |  |
| RMS<sup>f</sup> deviations |  |  |
| Bond lengths (Å) | 0.014 |  |
| Bond angles (°) | 1.393 |  |
| Ramachandran plot |  |  |
| % favored | 96.53 |  |
| % allowed | 2.48 |  |
| % outliers | 0.99 |  |
| PDB code | PDB: 6SWG |  |
| Diffraction data | DOI:15785/SBGRID/714 |  |

<sup>a</sup>Highest resolution shell is shown in parentheses.
<sup>b</sup> $R_{\text{sym}} = \sum_{hkl} \sum_i |I_{hkl,i} - \langle I \rangle_{hkl}| / \sum_{hkl} \sum_i I_{hkl,i}$ , where $I_{hkl}$ is the intensity of a reflection and $\langle I \rangle_{hkl}$ is the average of all observations of the reflection.
<sup>c</sup> $R_{\text{pim}} = \sum_{hkl} (N_{hkl} - 1)^{-1/2} \times \sum_i |I_{hkl,i} - \langle I \rangle_{hkl}| / \sum_{hkl} \sum_i I_{hkl,i}$ , where $I_{hkl}$ is the intensity of a reflection and $\langle I \rangle_{hkl}$ is the average of all observations of the reflection.
<sup>d</sup> $R_{\text{free}}$ , $R_{\text{work}}$ with 5% of $F_{\text{obs}}$ sequestered before refinement.
<sup>e</sup>Residual B-factors after TLS refinement. See PDB entry for TLS refinement parameters.
<sup>f</sup>R.M.S., root mean square.

To determine whether the binding interfaces observed in the Periphilin-TASOR complex are required for HUSH function, we used our structure to design point mutations in Periphilin predicted to interfere with Periphilin-TASOR complex assembly and measured the silencing activity of the mutants in the GFP reporter assay described above. Variants L326A and L333A/I337A were generated to target the Periphilin dimer interface; Periphilin L356R was generated to target both Periphilin-TASOR interfaces (**Fig. 2A**). The L356R variant failed to rescue reporter repression in Periphilin KO cells, whereas the L326A and L333A/I337A variants each had approximately half of the repression activity of wild-type Periphilin (**Fig. 2C**). Immunofluorescence microscopy with an anti-Periphilin antibody confirmed that all variants were expressed with the same nuclear localization as wild-type Periphilin (**Fig. 2D**). We note that the L356R variant that lacked silencing activity was expressed at higher levels than the partially active or wild-type variants. We conclude that the leucine zipper interactions between Periphilin and TASOR are absolutely required for HUSH function and that the fully active HUSH complex contains a Periphilin homodimer and a single TASOR molecule (or multiple 2:1 Periphilin:TASOR heterotrimers if additional oligomerization interfaces are present elsewhere in the HUSH complex).

### Redundancy within the Periphilin N-terminal region required for HUSH function

The requirement of Periphilin residues 1-127 for HUSH-dependent silencing but not HUSH complex assembly (**Fig. 1**) raises the question of how this N-terminal domain (NTD) contributes to HUSH function. Residues 20-291 of Periphilin are predicted to be unstructured (**Fig. 3A**). The sequence is more polar than hydrophobic, with clusters of alternating positive and negative net charge but an approximately neutral overall net charge. The NTD of Periphilin has a greater than average number of serine, arginine, tyrosine and negatively charged residues (see **Fig. 5A** below). Residues 147-222 (140-215 in isoform 1) are a serine-rich domain with six candidate serine phosphorylation sites, and a further three candidate sites at nearby residues 117, 121 and 140 (Olsen *et al*, 2006; Olsen *et al*, 2010; Zhou *et al*, 2013). To shed light on the role of these elements in HUSH-dependent silencing we generated Periphilin variants with various deletions in the NTD and measured their reporter repression activity over 21 days. Consistent with the reporter repression data shown in **Fig. 1A**, the 1Δ70 mutant repressed reporter expression to the same extent and at the same rate as wild-type Periphilin, whereas the 1Δ127 variant had no repression activity (**Fig. 3B**). Unexpectedly, however, addition of residues 1-70 to the 1Δ127 variant restored repression activity to 70% of wild-type activity. Hence, deletion of Periphilin residues 1-70 does not affect HUSH activity but these residues restore activity if residues 71-126 are deleted. Immunofluorescence microscopy confirmed that all variants were expressed in the nucleus, at similar levels (**Fig. 3C**). We conclude that residues 1-70 and 71-126 contribute to HUSH-dependent silencing in a redundant manner. Together, the amino acid sequence and redundant activities of the Periphilin NTD suggest that it is intrinsically disordered and hence that its contribution to HUSH activity stems from primary sequence attributes rather than tertiary structure.

**Fig. 3.**
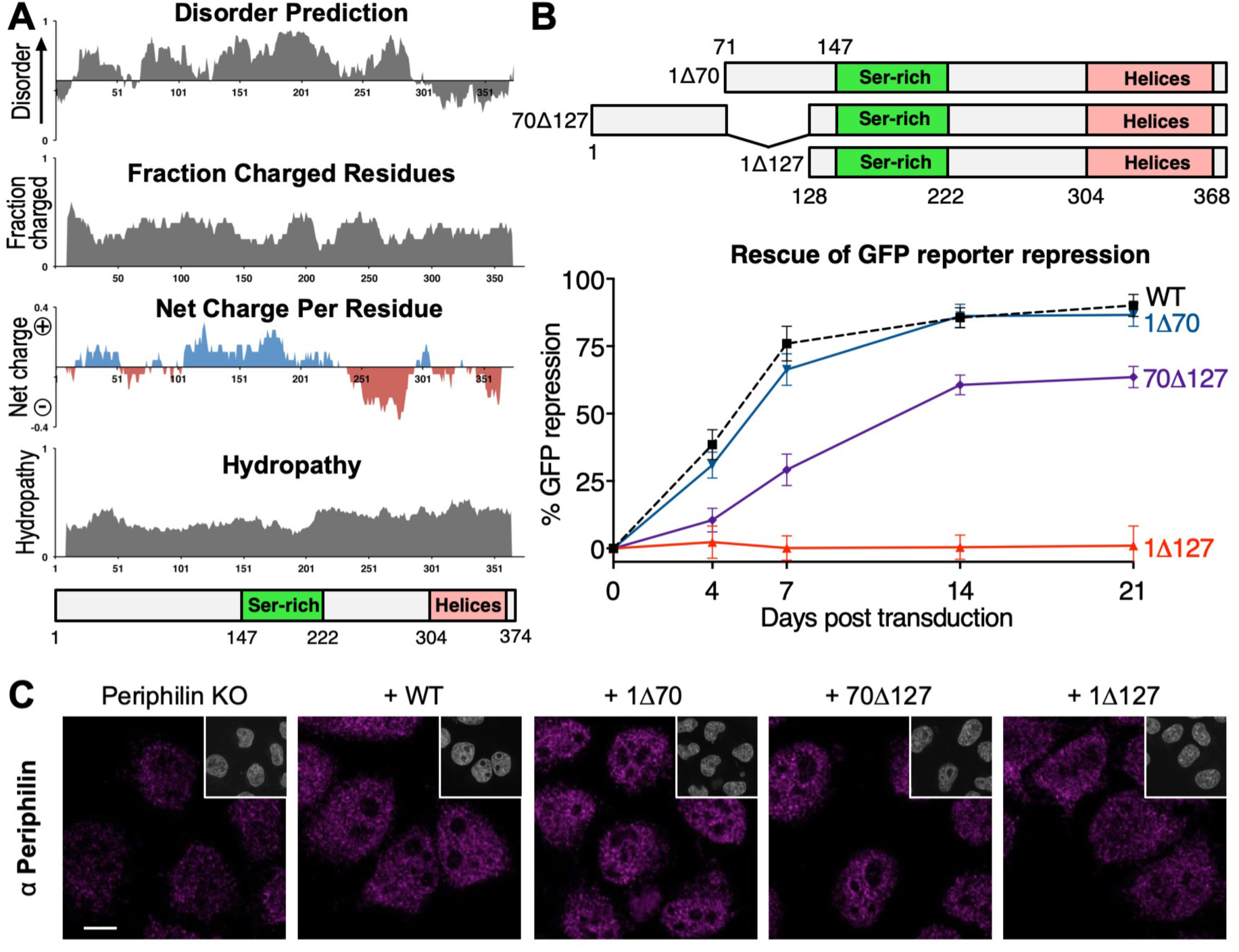
The NTD of Periphilin required for HUSH function contains partially redundant sequences predicted to be unstructured. (**A**) Predicted structural disorder, charge and hydropathy of Periphilin calculated with localCIDER (Holehouse *et al*, 2017). (**B**) Repression of a lentiviral GFP reporter in Periphilin KO cells complemented with Periphilin variants containing deletions in the NTD. Repression activity is calculated as in Fig. 2C. The WT curve (dotted line) is shared with contemporaneous experiments reported in Fig. 2C. (**C**) Immunofluorescence microscopy of Periphilin KO cells transduced with Periphilin N-terminal deletion variants. Cells were fixed 4 days post-transduction and stained with anti-Periphilin antibody (magenta) and DAPI (grey, insets). Scale bar, 10 µm.

**Fig. 4.**
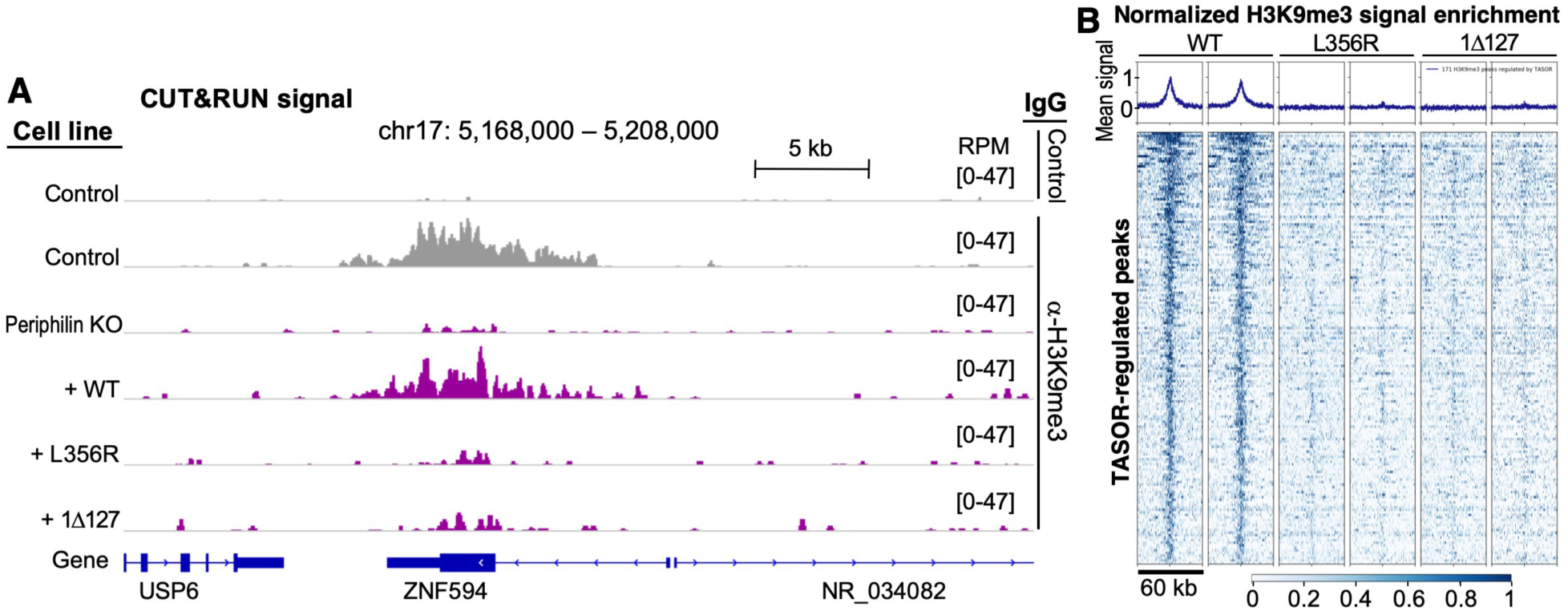
CUT&RUN genome-wide analysis of Periphilin and H3K9me3 distribution with wild-type and functionally deficient variants of Periphilin. (**A**) Snapshot of H3K9me3 distribution along the genome in the presence of different Periphilin variants. H3K9me3 distribution is shown at the ZNF594 locus, shown previously to be transcriptionally repressed by HUSH (Tchasovnikarova *et al*, 2015). Other representative snapshots are shown in **Fig. EV3**. An H3K9me3 track from wild-type HeLa cells (Control) and a track with a non-cognate IgG are shown in grey as positive and negative controls, respectively. The Periphilin-complemented tracks are in purple. All tracks were run in duplicate with similar results. RPM, reads per million, scaled to the total number of reads. (**B**) Heatmap showing CUT&RUN signal enrichment (normalized signal from complementing Periphilin construct minus normalized signal from Periphilin KO) over the 171 highest-confidence TASOR-regulated H3K9me3 peaks in the genome (Douse *et al*, 2019), centered on each peak, with a ±30 kb window. Both replicates are shown for WT, L356R and 1Δ127 Periphilin variants. The mean binned signal is shown above each heatmap.

**Fig. 5.**
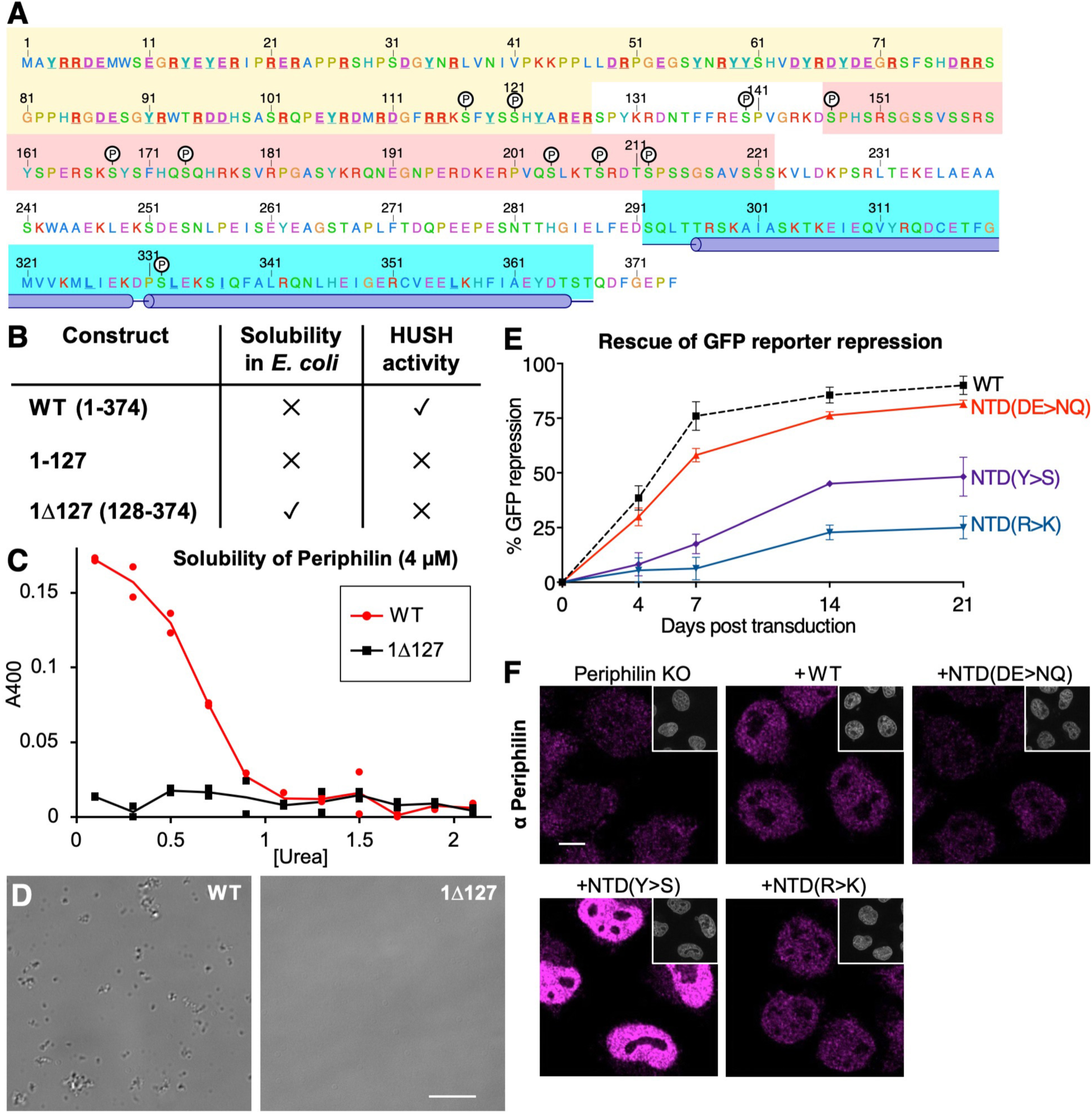
Physicochemical properties of the Periphilin NTD and contribution of enriched residues to HUSH activity. (**A**) Amino acid sequence of Periphilin-1 isoform 2 (UniProt Q8NEY8-2). Background shading color code: yellow, NTD; pink, Ser-rich region; blue, crystallized TASOR-binding region, with positions of the *α*-helices indicated. Candidate phosphorylation sites are labeled (P, black circles). Residues mutated in this study are underlined and in bold typeface. (**B**) Solubility of wild-type and terminal-deletion variants of Periphilin expressed in *E. coli*. The presence of HUSH-dependent silencing activity is indicated for each variant (see Fig. 3B). (**C**) Solubility of WT and 1Δ127 Periphilin variants, measured as absorbance at 400 nm (A400) due to light scattering, as a function of urea concentration in the buffer. Representative of two independent experiments. (**D**) Differential interference contrast (DIC) microscopy of WT and 1Δ127 Periphilin variants (4 µM) in buffer containing 0.5 M urea. Scale bar, 10 µm. (**E**) Repression of a lentiviral GFP reporter in Periphilin KO cells complemented with Periphilin variants with all Asp/Glu, Arg or Tyr in the NTD mutated to Asn/Gln, NTD(DE>NQ); Lys NTD(R>K); or Ser NTD(Y>S), respectively. Repression activity is calculated as above. The WT curve (dotted line) is shared with contemporaneous experiments reported in Fig. 2C. (**F**) Immunofluorescence microscopy of Periphilin KO cells transduced with Periphilin variants NTD(DE>NQ), NTD(R>K) or NTD(Y>S). Cells were fixed 4 days post-transduction and stained with anti-Periphilin antibody (magenta) and DAPI (grey, insets). Scale bar, 10 µm.

### The NTD and TASOR-binding site are required for H3K9 methylation at HUSH-regulated loci

Deposition of the repressive epigenetic mark H3K9me3 by SETDB1 is an essential component of HUSH-dependent silencing (Tchasovnikarova *et al*, 2015). To assess the importance of the Periphilin N- and C-terminal regions in H3K9 trimethylation, we measured the genome-wide distribution of H3K9me3 in cells expressing different Periphilin variants with the CUT&RUN (Cleavage Under Targets and Release Using Nuclease) epigenomic profiling method (Skene & Henikoff, 2017). We found that in Periphilin KO cells H3K9 methylation was lost or markedly reduced at hundreds of loci (**Figs. 4** and **EV3**). This pattern of H3K9me3 loss recapitulates that seen in previous ChIP-seq (Chromatin immunoprecipitation followed by sequencing) studies on cells in which TASOR, MPP8 or Periphilin were knocked out (Douse *et al*, 2019; Liu *et al*, 2018; Tchasovnikarova *et al*, 2015). Complementation of Periphilin KO cells with the L356R TASOR-binding mutant or with the 1Δ127 NTD deletion mutant failed to restore H3K9 methylation at HUSH-regulated loci (**Figs. 4** and **EV3**). The similar genome-wide distribution of H3K9me3 in Periphilin KO, L356R and 1Δ127 cell lines indicates that both the disordered NTD and the folded TASOR-binding domain of Periphilin are required for HUSH-dependent H3K9 methylation.

### Arginine and tyrosine residues in the NTD contribute to HUSH function

The physicochemical properties of the Periphilin NTD are reminiscent of the properties that govern the self-assembly of proteins into biomolecular condensates, in particular the Fused in Sarcoma (FUS) family of RNA-binding scaffold proteins. FUS family proteins contain N- and C-terminal intrinsically disordered regions with low sequence complexity resulting from a preponderance of specific subsets of amino acids (Malinovska *et al*, 2013). The N-terminal disordered region, known as the prion-like domain for its genetic association with prion-like inheritance in yeast and age-related neurodegenerative diseases in humans, is enriched in serine, glycine, tyrosine, glutamine, asparagine and proline (Alberti *et al*, 2009; Malinovska *et al*, 2013). The C-terminal region comprises one or more folded RNA recognition motifs (RRMs) interspersed with low-complexity sequences enriched in arginine and glycine. Arginine-tyrosine interactions and *π*-stacking of tyrosine-containing strands into kinked *β*-sheet fibrils in these disordered regions can non-covalently crosslink the polypeptide chains into liquid- or gel-like condensates, which manifest in the cell as phase separations or membraneless compartments (Hughes *et al*, 2018; Kato *et al*, 2012; Schwartz *et al*, 2013; Wang *et al*, 2018). The arginine-tyrosine interactions that promote phase separation of FUS family proteins are stabilized by complementary negative electrostatic charges from aspartate and glutamate residues in the prion-like domain (Wang *et al*, 2018). An excess of negative charge in FUS from multiple serine phosphorylation (or phosphomimetic mutations) decreases phase separation (Monahan *et al*, 2017). The NTD of Periphilin contains a similar sequence bias as the disordered C-terminal regions of FUS family proteins, with a marked enrichment of serine, arginine, tyrosine, aspartate and glutamate residues (**Figs. 5A, 6A**). Moreover, Periphilin has approximately the same number of positively and negatively charged residues, and has 10 potential serine phosphorylation sites, a similar number as FUS.

**Fig. 6.**
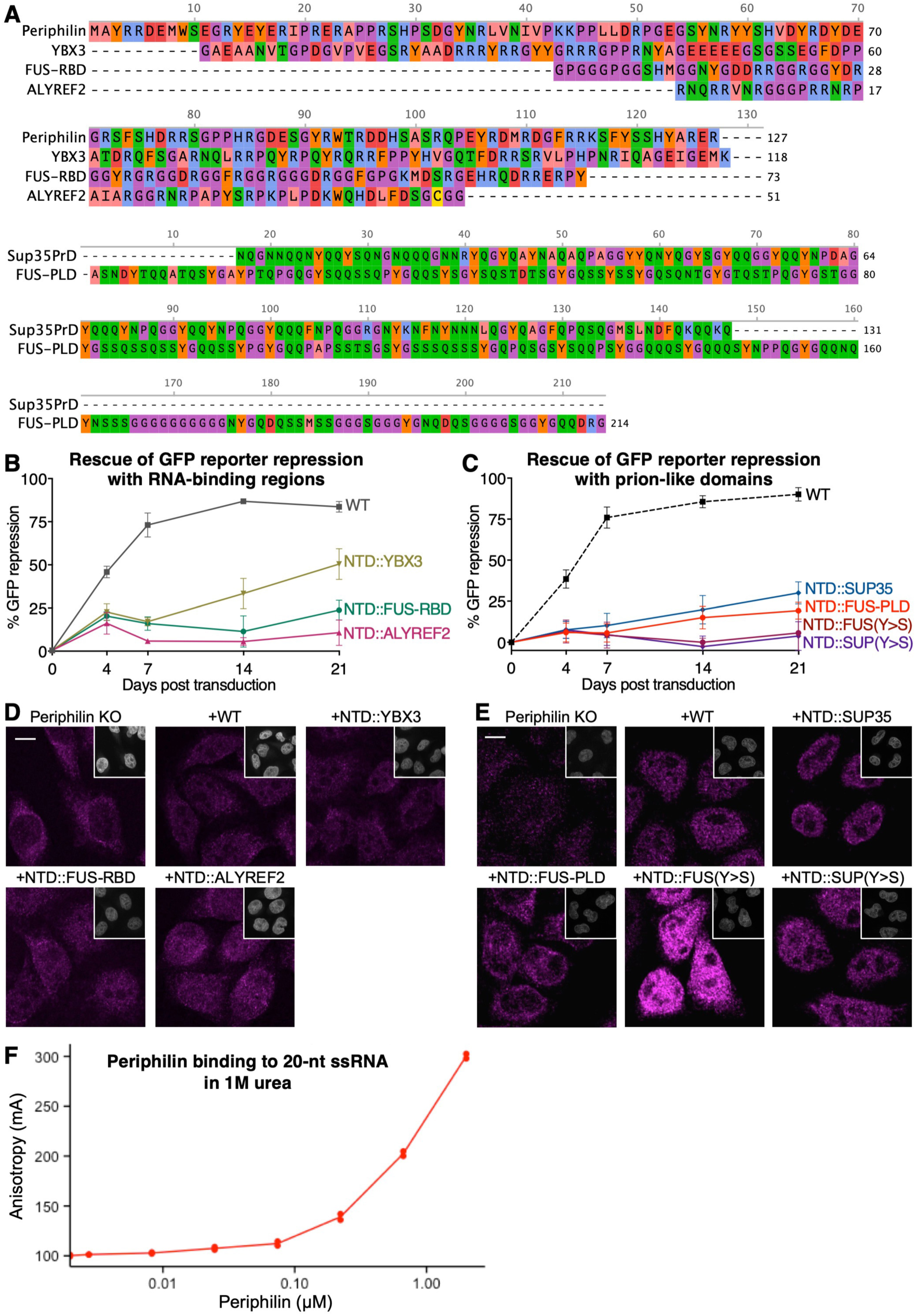
Complementation of NTD deletion with disordered regions from RNA-binding or prion-forming polypeptides partially rescues HUSH function. (**A**) Sequences used in this study to functionally complement the NTD of Periphilin in the 1Δ127 variant: YBX3, human Y-box-binding protein 3 residues 151-268, UniProt P16989-1; SUP35, *Saccharomyces cerevisiae* SUP35 prion domain, UniProt P05453 residues 5-135; FUS-RBD, human Fused in Sarcoma disordered RNA-binding region, UniProt P35637 residues 454-526; FUS-PLD, human Fused in Sarcoma prion-like low complexity domain, UniProt P35637 residues 2-214; ALYREF2, mouse Aly/RNA export factor 2 residues 17-67, UniProt Q9JJW6.1. (**B**) and (**C**) Repression of a lentiviral GFP reporter in Periphilin KO cells complemented with Periphilin variants with the NTD (residues 1-127) replaced by: (**B**) the disordered RNA-binding regions from ALYREF2, YBX3 or FUS; (**C**) the prion-like domain from FUS or the prion domain of SUP35, wild-type or with all tyrosine residues mutated to serine (Y>S). Residue ranges and sequence accession numbers are listed in (**A**). Repression activity is calculated as above. The WT curve in (**C**) is shared with contemporaneous experiments reported in Fig. 2C and is shown here as a dotted line. The WT curve in (**B**) is part of a separate experiment and is represented as a solid line. (**D**) and (**E**) Immunofluorescence microscopy of Periphilin KO cells transduced with the Periphilin variants shown in (**B**) and (**C**), respectively. Cells were fixed 4 days post-transduction and stained with anti-Periphilin antibody (magenta) and DAPI (grey, insets). (**F**) Binding of recombinant Periphilin to a 20-nucleotide single-stranded RNA (20-nt ssRNA) labeled at the 5’ end with 6-carboxyfluorescein (6-FAM). Periphilin was titrated into a solution of 20 nM fluorescein-labeled ssRNA, in the presence of 1 M urea to minimize Periphilin aggregation. ssRNA binding was measured as an increase in fluorescence polarization anisotropy.

To assess the potential of Periphilin to form condensates we expressed various Periphilin recombinant protein constructs in *E. coli.* Full-length Periphilin and a construct spanning the NTD alone (residues 1-127) were both insoluble and could not be purified from cell lysates under native conditions (**Fig. 5B**). The 1Δ127 variant lacking the NTD was soluble but lacked repression activity as noted above. In contrast to FUS family proteins, which undergo phase separation and form hydrogels reversibly at low salt concentrations, Periphilin constructs containing the NTD remained insoluble even at higher than physiological salt concentrations (0.3 M NaCl). Full-length Periphilin could be solubilized and purified in the presence of 8 M urea but upon dilution of the urea to below 1 M the protein came out of solution, reversibly, forming solid aggregates detectable by absorbance in the visible light spectrum and by differential interference contrast microscopy (**Fig. 5C,D**). Hence, the NTD, which is required for HUSH activity, induces Periphilin to self-aggregate without undergoing phase separation or hydrogel formation as seen in FUS family proteins.

Among the amino acids enriched in the disordered regions of Periphilin and FUS family proteins, tyrosine and arginine residues govern the phase separation properties of FUS family proteins (Wang *et al*, 2018). Mutation of tyrosine residues to serine, or arginine to alanine, in FUS disordered regions diminishes or abrogates phase separation and hydrogel formation (Kato *et al*, 2012; Wang *et al*, 2018). To determine whether arginine and tyrosine residues contribute to Periphilin self-aggregation, we generated Periphilin variants with all 24 arginine residues in the NTD mutated to lysine, NTD(R>K), or with all 13 tyrosine residues mutated to serine, NTD(Y>S) and measured the silencing activity of the mutants in our GFP reporter assay. Both variants reduced HUSH-dependent repression (**Fig. 5E**). The NTD(R>K) variant, despite retaining the net charge of wild-type Periphilin, had less than 10% of wild-type activity 7 days post-transduction, and approximately one quarter of wild-type activity after 21 days. The NTD(Y>S) variant appeared less impaired, with 20% of wild-type activity after 7 days and 50% after 21 days. However, the NTD(Y>S) variant was expressed at significantly higher levels than the NTD(R>K) and wild-type variants, suggesting that the impairment of the NTD(R>K) and NTD(Y>S) variants would be comparable at identical expression levels (**Fig. 5F**).

Arginine-tyrosine interactions have been proposed to be stabilized by negatively charged residues in FUS family proteins. Mutation of negatively charged residues in FUS family proteins decreases their overall phase separation potential (Wang *et al*, 2018). In contrast, the negatively charged residues in the NTD were not required for silencing. Indeed, Periphilin variant NTD(DE>NQ) with all 22 aspartate or glutamate residues mutated to asparagine or glutamine, respectively, had repression activity similar to wild-type (**Fig. 5E**). Immunofluorescence microscopy confirmed the DE>NQ, R>K and Y>S variants had nuclear localization, although the latter variant was more abundantly expressed (**Fig. 5F**).

### Disordered polypeptides with self-associating or RNA-binding properties partially complement NTD deletion

The similarity of the amino acid sequence bias in the Periphilin NTD and the disordered regions of RNA-binding domains from FUS family proteins raises the question of whether these low-complexity sequences have similar biophysical properties, which in Periphilin contribute directly to HUSH-dependent silencing. Insertion of the disordered portion of the RNA-binding domain from FUS (**Fig. 6A**) into the 1Δ127 variant restored HUSH repression activity to approximately 25% of wild-type (**Fig. 6B**, NTD::FUS-RBD). To determine whether self-association of a disordered polypeptide *per se* is sufficient to support HUSH silencing we generated variants containing the prion domain from yeast SUP35, or the prion-like domain of FUS in place of the NTD (NTD::SUP35 and NTD::FUS-PLD, respectively). Prion domains have a different sequence bias: enrichment of glutamine, asparagine and tyrosine and depletion of charged residues (**Fig. 6A**) (Alberti *et al*, 2009). Prion domains form highly stable steric zipper-type amyloid fibers distinct from the reversible associative polymers formed by FUS (Hughes *et al*, 2018; Murray *et al*, 2017).

Nevertheless, the NTD::SUP35 and NTD::FUS-PLD variants restored repression activity to 30% and 20% of wild-type (**Fig. 6C**), respectively, suggesting prion-like aggregation partially functionally complements the NTD, to a comparable extent as the arginine/glycine-rich region of FUS.

Tyrosine residues are essential for the aggregation of FUS and the amyloidogenic properties of prion proteins (Alexandrov *et al*, 2008; Kato *et al*, 2012; Murray *et al*, 2017; Wang *et al*, 2018). Mutation of all tyrosine residues to serine in the complementing sequences of FUS-PLD and SUP35 abrogated their silencing activity (**Fig. 6C**, NTD::FUS(Y>S) and NTD::SUP(Y>S)).

A large proportion of proteins with low-complexity disordered regions bind RNA via arginine-rich sequences. Many of these RNA-binding proteins self-associate into liquid or gel phases, or form amyloid (or amyloid-like) fibers (Jarvelin *et al*, 2016; King *et al*, 2012). The same arginine/glycine-rich regions of FUS family proteins that mediate phase separation also bind RNA, and RNA binding nucleates higher-order assembly of FUS (Schwartz *et al*, 2013). Moreover, prion proteins are strongly enriched for RNA-binding proteins (Alberti *et al*, 2009). Formation of ribonucleoprotein complexes (RNPs)— in particular with mRNA— through phase separation or fibril formation is emerging as a central mechanism of co- and post-transcriptional regulation (Jarvelin *et al*, 2016). To assess the potential contribution of RNA binding by Periphilin to silencing, we replaced the NTD with RNA-binding polypeptides from two RNA-binding proteins (**Fig. 6A**), Y-box-binding protein 3 (YBX3) and Aly/RNA export factor 2 (ALYREF2). YBX3 a member of the cold shock domain (CSD) protein family that binds mRNA without sequence specificity via a disordered C-terminal tail rich in aromatic, basic and phosphorylated residues (Manival *et al*, 2001; Matsumoto *et al*, 1996). YBX3 was recently shown to repress translation of certain mRNAs (Cooke *et al*, 2019). ALYREF2 contributes to mRNA export by packaging mRNA into RNPs through interactions with an arginine-rich disordered N-terminal tail. The NTD::YBX3 variant restored HUSH repression activity to the greatest extent of any of the complementing sequences tested, with 50% of wild-type Periphilin 21 days post-transduction (**Fig. 6B**). In contrast, the NTD::ALYREF2 variant did not restore repression. Immunofluorescence microscopy confirmed all variants were expressed with nuclear localization, although the NTD::FUS(Y>S) variant was expressed at higher levels than the other variants (**Fig. 6D,E**). We note that the complementing YBX3 sequence and the Periphilin NTD both have a greater number of alternating positively and negatively charged residues than the other complementing sequences (**Fig. 6A**).

Efforts to measure RNA binding by Periphilin were hampered by the insolubility of purified protein constructs containing the NTD. In the presence of 1 M urea to minimize aggregation, binding was observed between full-length Periphilin and a 20-nucleotide single-stranded RNA labeled with fluorescein, as judged from an increase in fluorescence polarization anisotropy (**Fig. 6F**). The anisotropy signal did not saturate, consistent with the interpretation that Periphilin bound to the oligonucleotide non-specifically through multiple interaction sites.

## Discussion

We have identified the key structural and biochemical properties of Periphilin necessary for epigenetic silencing by the HUSH complex. The C-terminal coiled-coil domain directs HUSH complex assembly by dimerizing and binding TASOR through *α*-helical coiled-coil interactions. How the N-terminal region (NTD) of Periphilin contributes to silencing is more difficult to pinpoint due to its intrinsic structural disorder and its propensity to aggregate. We note that self-aggregation of the NTD correlates with HUSH function, as truncations that inhibit aggregation also inhibit silencing. As in self-associating disordered regions from many other proteins including FUS-family proteins, the NTD of Periphilin is enriched in tyrosine and arginine residues. These residues are required for HUSH repression activity. Arginine-tyrosine *π*-stacking interactions are essential for the aggregation of FUS, and tyrosine residues contribute to the amyloidogenic properties of prion proteins. Hence, NTD self-association through arginine-tyrosine *π*-stacking interactions could play a role in HUSH silencing. Consistent with this notion, lysine failed to functionally substitute for arginine in silencing assays with our 1NTD(R>K) Periphilin variant.

Alongside the broad similarities between the Periphilin NTD and disordered regions from other arginine/tyrosine-rich proteins, the NTD has certain distinguishing properties. The sequence complexity of the NTD is not as low as in the disordered regions that drive phase separation of FUS family proteins. Periphilin lacks tyrosine residues flanked on both sides by serine or glycine to form [G/S][Y/F][G/S] motifs, which are hallmarks of fiber-forming Low-complexity Aromatic-Rich Kinked Segments (LARKS). Periphilin also contains a greater proportion of charged residues than FUS-family and prion proteins and may acquire further negative charges through phosphorylation. Moreover, the NTD induces the formation of solid aggregates rather than liquid-like phases, hydrogels or fibers. These distinguishing features may explain why the NTD was not fully complemented by the disordered regions from FUS or Sup35 in our HUSH silencing assays.

On balance, the similarities between the NTD and self-associating disordered regions from FUS-family proteins outweigh the differences. Indeed, counterbalancing the differences listed above, the NTD does contain sequences that resemble LARKS or are predicted to have amyloid-forming potential, for example SFYSSHYA, with a stacking free energy of −27 kcal/mol predicted by ZipperDB (Goldschmidt *et al*, 2010). Second, the negatively charged residues in the NTD, though more abundant than in FUS and Sup35, are not essential for HUSH function. Furthermore, membraneless compartments formed by biomolecular condensates have been reported previously to have the characteristics of a solid rather than a liquid or gel (Banani *et al*, 2017; Shin & Brangwynne, 2017). Hence, the most plausible mechanism for Periphilin NTD aggregation is via arginine-tyrosine *π*-stacking interactions, like FUS-family proteins but with greater cooperativity, resulting in a more abrupt transition from the soluble state to a solid aggregated state. Whether and how self-association via this mechanism translates into silencing activity in the HUSH complex remains unclear.

The biochemical properties of the NTD suggest that one of its functions in HUSH-dependent silencing may be to bind RNA. A majority of proteins with arginine-rich disordered regions bind RNA and self-associate into ribonucleoprotein (RNP) fibers or condensates (Jarvelin *et al*, 2016; King *et al*, 2012). Polymerization of proteins— such as proteins from cold shock domain (CSD) family— on mRNA is emerging as a central mechanism to repress protein expression co- or post-transcriptionally (Jarvelin *et al*, 2016). Notably, the disordered RNA-binding region of CSD-family protein YBX3, which binds certain mRNAs and represses their translation (Cooke *et al*, 2019), functionally complemented the Periphilin NTD to a greater extent than any of the other sequences we tested. As in other CSD proteins, the disordered RNA-binding region of YBX3 is enriched in aromatic, basic and phosphorylated residues and binds mRNA without sequence specificity (Manival *et al*, 2001; Matsumoto *et al*, 1996). Whether Periphilin forms RNPs with mRNA remains unknown, but we found that Periphilin bound a single-stranded RNA oligonucleotide in 1 M urea (in the absence of urea, light scattering from Periphilin aggregates precluded ligand binding measurements). Additionally, the approximately linear shape of the binding curve was consistent with non-specific binding via multiple interaction sites. Binding of Periphilin to mRNA could explain the known propensity of HUSH to silence genes that are actively being transcribed (Douse *et al*, 2019; Liu *et al*, 2018). In support of HUSH binding to nascent transcripts, Periphilin, TASOR (as C3orf63) and Periphilin mRNA were identified in a proteomic screen for protein-mRNA interactions in HeLa cells (Castello *et al*, 2012). Moreover, artificially increasing transcription of a transgene increases recruitment of HUSH to that locus (Liu *et al*, 2018). Why the HUSH complex preferentially binds to intronic LINE-1 elements within actively transcribed genes with some degree of sequence specificity (Liu *et al*, 2018), remains to be determined. We note that the cold shock domains of YBX3 and other Y-box binding proteins bind to specific sequences or structures in the untranslated regions of their target mRNAs (Cooke *et al*, 2019; Yang *et al*, 2019). Although the HUSH complex does not contain any classical RNA recognition motifs or cold shock domains, the possibility that HUSH contains a motif conferring specificity for RNA sequence or structure cannot be excluded.

The work presented and discussed here prompts us to propose the following model for how Periphilin may function in silencing. The NTD of Periphilin may bind nascent transcripts, with multiple Periphilin molecules binding to each target mRNA. The self-aggregation properties of NTD could then lead to the formation of large mRNPs. Transcripts within these mRNPs would be less accessible to transcription or mRNA processing machinery, thereby repressing expression. Other HUSH components or effectors, tethered via the C-terminal domain of Periphilin, could then sense and modify the epigenetic landscape or chromatin structure at the target site. This would include deposition of the transcriptionally repressive H3K9me3 mark. Further studies will be necessary to test and refine this model. This work provides a foundation to design new epigenetic therapies targeting HUSH to treat autoimmune diseases, cancer and retroviral infections.

## Materials and Methods

### Protein sequence analysis

We used CIDER software developed for the analysis of intrinsically disordered proteins (Holehouse *et al*, 2017) to extract Periphilin sequence parameters including predicted structural disorder, charge and hydropathy.

### Protein expression vectors

Synthetic genes encoding codon-optimized Periphilin-1 isoform 2 (UniProt Q8NEY8-2, residues 285-374) and TASOR isoform 1 (UniProt Q9UK61-1, residues 1014-1095) were cloned sequentially into pRSF-Duet vector (Novagen) using HiFi Assembly (NEB), producing an N-terminally hexahistidine (His_6_) tagged Periphilin and untagged TASOR fragments. These constructs were used to produce Periphilin-TASOR complex for biophysical characterization by NMR and native mass spectrometry (see below). Following identification of disordered regions by NMR, a shorter Periphilin variant (residues 292-367) was cloned, preceded by a His_6_ tag and a TEV protease cleavage site (ENLYFQG). This construct was used for crystallization and SEC-MALS of the Periphilin-TASOR complex. Full-length, 1-127, and 128-374 Periphilin codon-optimized for *Escherichia coli* were cloned into pET15b plasmid (Novagen), with N-terminal His_6_tags followed by a thrombin protease cleavage site (LVPRGS); these constructs were used for biophysical characterization of the Periphilin N-terminus.

### Lentivirus complementation assay vectors

Wild-type Periphilin was cloned into lentiviral vector pHRSIN-PSFFV-V5-Periphilin-PPGK-Hygro as described (Tchasovnikarova *et al*, 2015). For Periphilin mutants L356R, L326A and L333A/I337A, the vector was digested with MluI and NotI to remove the residues 295-374 of the insert. Synthetic dsDNA fragments carrying the mutations were inserted into the gel-purified vector by HiFi Assembly. For the 1Δ127, 1Δ70, and 70Δ127 deletion mutants, the vector was digested with KpnI and NotI to remove the Periphilin sequence and one or two PCR fragments corresponding to the retained sequences were inserted by assembly. The NTD(DE>NQ), NTD(R>K) and NTD(Y>S) variants were generated by assembling a synthetic dsDNA encoding Periphilin residues 1-127 with the desired mutations with a PCR product encoding Periphilin residues 128-374 and KpnI/NotI-digested vector. Constructs with complementing sequences from other proteins were generated in the same manner but using a synthetic dsDNA encoding: FUS PLD (UniProt P35637, residues 2-214); SUP35 PrD (UniProt P05453, residues 5-135); ALYREF2 residues 17-67 (UniProt Q9JJW6.1), YBX3 residues 151-268 (UniProt P16989-1), and FUS RBD (UniProt P35637, residues 454-526).

### Cell lines and lentivirus production

Periphilin knockout HeLa cells carrying integrated GFP reporter and HEK 293T were maintained in RPMI supplemented with 10% fetal calf serum, 50 U/ml penicillin, 50 µg/ml streptomycin. Lentiviruses were produced by cotransfection of HEK 293T cells at 95% confluence in 6-well plates with 3 µg of the following plasmids in 1:2:2 molar ratio: pMD2.G carrying glycoprotein VSV-G, pCMVΔ8.91 carrying replicative genes, and the pHRSIN-based lentiviral backbone containing hygromycin resistance and Periphilin. The plasmids were mixed with 200 µl serum-free medium, and 15 µl PEI, incubated at room temperature for 20 min and applied to cells. Media was exchanged 4 h post-transfection and supernatants containing lentiviruses were harvested 48 h post-transfection by filtering through a 0.45 µm filter and stored at −80°C.

### Reporter silencing assay

Reporter silencing activity of WT and mutant Periphilin was measured by infecting the HeLa reporter cell line with lentiviruses carrying Periphilin variants and monitoring GFP fluorescence for 21 days post-transduction (Tchasovnikarova *et al*, 2015). Periphilin KO HeLa cells in 24-well plates were overlaid with 150 µl of lentiviral supernatants and 8 µg/ml polybrene and centrifuged for 90 min at room temperature at 1000 g. After 24 h incubation cells were trypsinized and seeded into flasks with selection media containing 400 µg/ml hygromycin. Fresh media was added every other day. Hygromycin was removed from the media after 7 days in culture. For flow cytometry cells were trypsinized, washed in PBS, counted, and resuspended at 1×10^6^ cells per ml in PBS supplemented with 2% fetal calf serum. GFP fluorescence was recorded with an Eclipse flow cytometer (iCYT) from >1×10^5^ cells per sample. The cells were gated on live single-cell population based on forward and side scatter in FlowJo (BD Life Sciences). The geometric mean of the GFP fluorescence of the whole live population was determined without further gating and values exported to Excel (see **Data Set EV1**). Since gene expression data is log-normal, we converted GFP fluorescence to percent repression with the formula: %GFP Repression = log_10_(GeoMeanPopulation)*m + a, where m = 100% / [log_10_(GeoMeanWT) − log10(GeoMeanKO)] and b = −m*log10(GeoMeanKO). This transformation assigned the Periphilin KO population a value of 0% repression and WT HeLa cells 100% repression. The values of m and b used were −59.7 and 196.1, respectively, for all experiments except those in **Fig. 6C**, where they were −59.5 and 210.0. The difference in vertical offset was due to a laser upgrade on the instrument.

### Co-immunoprecipitation and Western blotting

For co-immunoprecipitation, cells were lysed in 1% NP-40 in TBS plus 10 mM iodoacetamide, 0.5 mM phenylmethylsulfonyl fluoride (PMSF) and benzonase (Sigma-Aldrich) for 30 min. Protein A and IgG-sepharose resin was added to the lysates along with primary antibody. The suspension was incubated for 2 h at 4°C and the resin was washed three times in lysis buffer. For Western blotting, cells were lysed with lysis buffer containing 1% SDS instead of 1% NP-40. For SDS-PAGE analysis, resins or lysates were heated to 70°C in SDS sample buffer for 10 min and run on a polyacrylamide gel. Gels were blotted onto PVDF membranes (Millipore). Blots were blocked in 5% milk in PBS, 0.2% Tween-20 and incubated with primary antibody diluted in blocking solution. As the Periphilin antibody was unable to detect its epitope under NP-40 lysis conditions, we used a mouse antibody against the V5 tag (Abcam, ab27671) as the primary antibody for Periphilin. For TASOR, the primary antibody was rabbit α-TASOR (Atlas, HPA006735). Blots were imaged with West Pico or West Dura (Thermo Fisher Scientific).

### Protein expression and purification

*E. coli* BL21 (DE3) cells (New England BioLabs) were transformed with pRSF-DUET constructs expressing TASOR-Periphilin complex and selected on kanamycin plates. For native protein expression, overnight cultures were diluted 1:200 into 2 l of LB. Cultures were induced with 100 mM IPTG at OD_600_ 0.6, incubated at 37°C for 2 h, harvested, resuspended in 50 ml Buffer A (20 mM HEPES pH 7.4, 0.5 M NaCl, 0.5 mM TCEP) and frozen in liquid nitrogen. The cells were thawed at room temperature, supplemented with 1 µL benzonase (Sigma) and lysed by sonication. The lysates were purified by centrifugation and filtering over a 0.45 µm filter. The clarified lysates were applied to 1-ml HisTrap columns (GE Healthcare), using one column per liter culture, washed with 20 column volumes of Buffer A and eluted in with 0.5 M imidazole in Buffer A. To purify non-cleavable His_6_-tagged Periphilin residues 285-374, the eluate was desalted into Buffer QA (20 mM NaCl, 20 mM HEPES pH 7.4, 0.5 mM TCEP) using a HiPrep desalting column (GE Healthcare), bound to MonoQ ion-exchange column (GE Healthcare) and eluted in a gradient of Buffer QA and BufferQB (1 M NaCl, 20 mM HEPES pH 7.4, 0.5 mM TCEP). Size-exclusion chromatography (SEC) on a Superdex 200 10/300 column (GE Healthcare) in PBS supplemented with 1mM TCEP and 0.05% sodium azide completed purification. For the N^15^ and N^15^/C^13^-labeled protein expression, the overnight starter culture was grown in complete unlabeled minimal medium and used to inoculate (1:100 v/v) 800 ml complete labeled media. Labeled proteins were purified as described above. For the construct expressing the TEV-cleavable His_6_-tagged Periphilin residues 392-367, Ni-affinity purification was followed by overnight digestion with TEV protease at 22°C, desalting into Buffer SA (20 mM NaCl, 20 mM Acetate pH 4.55, 0.5 mM TCEP), binding to a MonoS column, and elution against Buffer SB (1 M NaCl, 20 mM sodium acetate pH 4.55, 0.5 mM TCEP). SEC on a Superdex 200 column in 20 mM HEPES pH 7.4, 0.1 M NaCl, 0.5 mM TCEP completed purification. To purify full-length Periphilin, we followed the QIAexpressionist batch purification protocol under denaturing conditions (QIAGEN) followed by purification on a Superdex 200 column in 6 M urea, 0.2 M sodium phosphate pH 7.4, 20 mM TRIS pH 7.4, 1 mM TCEP.

### X-ray crystallography

Crystals were grown at 18°C by sitting drop vapor diffusion. Purified Periphilin-TASOR complex was mixed with an equal volume of reservoir solution: 0.1 M Citrate pH 4.5, 1 M ammonium sulfate. Crystals were harvested into a 70:30 mix of mother liquor to protein buffer supplemented with 20% DMSO with or without 1 M NaBr. Crystals were frozen in liquid nitrogen. X-ray diffraction data were collected at 100 K at Diamond Light Source (DLS) beamline I04-1. Automatic experimental phasing pipelines implemented at DLS including CRANK2 (Skubak & Pannu, 2013) determined phases with the single anomalous dispersion (SAD) method using bromine as the heavy atom. A polyalanine model built with CRANK2 was used as a molecular replacement search model for the native dataset (without NaBr) in PHENIX (Adams *et al*, 2010). The atomic model was built with COOT (Emsley & Cowtan, 2004) and iteratively refined with PHENIX (Adams *et al*, 2010) at 2.5 Å resolution. See **Table 1** for data collection and refinement statistics.

### Size-exclusion chromatography and multi-angle light scattering (SEC-MALS) analysis

100 µl of protein sample was subjected to SEC at 293 K using a Superdex 200 10/300 column (GE Healthcare) pre-equilibrated in PBS at a flow rate of 0.5 ml min^−1^. The SEC system was coupled to a multi-angle light scattering (MALS) module (DAWN-8+, Wyatt Technology). Molar masses of peaks in the elution profile were calculated from the light scattering and protein concentration, quantified using the differential refractive index of the peak assuming a dn/dc of 0.186, using ASTRA6 (Wyatt Technology).

### Immunofluorescence microscopy

Cells were grown on glass cover slips and then fixed with 4% formaldehyde in PBS for 15 min. Cells were permeabilized with 0.1% Triton X100 in PBS and then blocked with 5% BSA in PBS. Samples were stained with primary anti-Periphilin antibody (Atlas, HPA038902) at dilution 1:500 for 1 h and after washing with blocking buffer with secondary anti-rabbit AlexaFluor 568 antibody diluted 1/500 for 1 h. Cover slips were mounted on microscopy glasses with ProLong Gold anti-fade reagent with DAPI (Invitrogen). Imaging was performed using Nikon Ti microscope equipped with CSU-X1 spinning disc confocal head (Yokogawa) and with Zeiss 780 system.

### CUT&RUN H3K9me3 genome profiling

We followed the protocol detailed by Henikoff and colleagues (Skene & Henikoff, 2017). Briefly, 250,000 cells (per antibody/cell line combination) were washed twice (20 mM HEPES pH 7.5, 0.15 M NaCl, 0.5 mM spermidine, 1x Roche complete protease inhibitors) and attached to ConA-coated magnetic beads (Bangs Laboratories) pre-activated in binding buffer (20 mM HEPES pH 7.9, 10 mM KCl, 1 mM CaCl_2_, 1 mM MnCl_2_). Cells bound to the beads were resuspended in 50 µl buffer (20 mM HEPES pH 7.5, 0.15 M NaCl, 0.5 mM Spermidine, 1x Roche complete protease inhibitors, 0.02% w/v digitonin, 2 mM EDTA) containing primary antibody (1:100 dilution). Incubation proceeded at 4°C overnight with gentle shaking. Tubes were placed on a magnet stand to remove unbound antibody and washed three times with 1 ml digitonin buffer (20 mM HEPES pH 7.5, 0.15 M NaCl, 0.5 mM Spermidine, 1x Roche complete protease inhibitors, 0.02% digitonin). pA-MNase (35 ng per tube, a generous gift from Steve Henikoff) was added in 50 µl digitonin buffer and incubated with the bead-bound cells at 4°C for 1 h. Beads were washed twice, resuspended in 100 µl digitonin buffer and chilled to 0-2°C. Genome cleavage was stimulated by addition of 2 mM CaCl_2_ (final), briefly vortexed and incubated at 0°C for 30 min. The reaction was quenched by addition of 100 µl 2x stop buffer (0.35 M NaCl, 20 mM EDTA, 4 mM EGTA, 0.02% digitonin, 50 ng/µl glycogen, 50 ng/µl RNase A, 10 fg/µl yeast spike-in DNA (a generous gift from Steve Henikoff)) and vortexing. After 10 min incubation at 37°C to release genomic fragments, cells and beads were pelleted by centrifugation (16,000 g, 5 min, 4°C) and fragments from the supernatant purified with a Nucleospin PCR clean-up kit (Macherey-Nagel). Illumina sequencing libraries were prepared using the Hyperprep kit (KAPA) with unique dual-indexed adapters (KAPA), pooled and sequenced on a NovaSeq6000 instrument. Paired-end reads (2×150) were aligned to the human and yeast genomes (hg38 and R64-1-1 respectively) using Bowtie2 (--local –very-sensitive-local –no-mixed –no-discordant –phred33 −I 10 −X 700) and converted to bam files with samtools. Conversion to bedgraph format and normalization was performed with bedtools genomecov (-bg -scale), where the scale factor was the inverse of the number of reads mapping to the yeast spike-in genome. CUT&RUN experiments to assess H3K9me3 regulation by Periphilin variants were done in biological duplicate. Peaks defined as HUSH-regulated were reported elsewhere (Douse *et al*, 2019). Normalized bigwig files were generated (UCSC), displayed in IGV (Broad Institute) and heatmaps plotted with computeMatrix and plotHeatmap commands (deepTools). Figures were prepared in Inkscape.

### Periphilin solubility and RNA-binding assays

Denatured protein samples in Denaturing Buffer (6M urea, 0.2 M sodium phosphate pH 7.4, 20 mM TRIS pH 7.4, 1 mM TCEP) were concentrated to 120 µM protein on a 10-kDa cutoff centrifugal concentrator. Solutions with increasing concentrations of urea were prepared by diluting 1 µl Periphilin into 30 µl buffer. The absorbance at 400 nm (OD_400_) was measured in duplicate on a ClarioSTAR plate reader (BMG Labtech).

RNA binding was measured by fluorescence polarization anisotropy. Periphilin was titrated into Urea Buffer (1 M urea, 0.2 M sodium phosphate pH 7.4, 20 mM TRIS pH 7.4, 1 mM TCEP) containing 20 nM of a single-stranded RNA oligonucleotide with the sequence CCUGUAAUCCCAGCACUUUG labeled at the 5’ end with 6-carboxyfluorescein (6-FAM). Fluorescence anisotropy of 30-µl samples was measured in 384-well black, clear-bottomed plates (Corning) with a ClarioSTAR plate reader using 482/530 nm filters.

### Differential interference contrast (DIC) microscopy

WT and 1Δ127 Periphilin variants in Denaturing Buffer (see above) were diluted into 0.5 M Urea Buffer (0.5 M urea, 0.2 M sodium phosphate pH 7.4, 20 mM TRIS pH 7.4, 1 mM TCEP). After 1 h incubation, 5 µl of each sample was applied to a glass dish (Ibidi) and imaged on a Nikon Ti2 microscope with a 100X objective.

### Statistics

No statistical methods were used to predetermine sample size, experiments were not randomized, and the investigators were not blinded to experimental outcomes. Reporter silencing assays were performed at least three times in independent experiments. Repression activity data are represented as the mean ± standard error of the mean (s.e.m.), calculated with PRISM 8 (GraphPad), with three biological replicates for all experiments except WT with ten replicates.

### Data availability

The structure factors and atomic coordinates were deposited in the Protein Data Bank with code PDB: 6SWG. The original experimental X-ray diffraction images were deposited in the SBGrid Data Bank (data.SBGrid.org), with Data ID 714, DOI:10.15785/SBGRID/714.

## Acknowledgements

We thank Steven Henikoff for the generous gift of pA-MNase and yeast spike-in DNA for CUT&RUN experiments, and MRC-LMB Scientific Computing for support. Crystallographic data were collected on beamline I04-1 at Diamond Light Source (DLS). Access to DLS (proposal MX15916) was supported by the Wellcome Trust, MRC, and BBSRC. We thank members of the Modis lab for insightful discussions. This work was supported by Wellcome Trust Senior Research Fellowships 101908/Z/13/Z and 217191/Z/19/Z to Y.M., Wellcome Trust Principal Research Fellowship 101835/Z/13/Z to P.J.L., and a BBSRC Future Leader Fellowship BB/N011791/1 to C.H.D.

## Author Contributions

Conceptualization, D.M.P., A.A., C.H.D., I.A.T., R.T.T., P.J.L. and Y.M.; Methodology, D.M.P., A.A., C.H.D., I.A.T., R.T.T., S.M.V.F, S.M., P.J.L. and Y.M.; Investigation, D.M.P., A.A., C.H.D., I.A.T., R.T.T., L.E.F., S.M., S.O. and Y.M.; Validation – crystal structure, Y.M.; Writing – Original Draft, Y.M.; Writing – Review & Editing, D.M.P., C.H.D., P.J.L. and Y.M.; Visualization, D.M.P, A.A., C.H.D., I.A.T., R.T.T., S.M. and Y.M.; Supervision, P.J.L. and Y.M.; Project Administration, Y.M.; Funding Acquisition, P.J.L. and Y.M.

## Supplementary Information

### Supplementary Methods

#### Nuclear magnetic resonance (NMR) spectroscopy

A ^1^H,^15^N BEST-TROSY NMR spectrum of the Periphilin-TASOR complex (^15^N,^13^C-labeled Periphilin residues 285-374 with an N-terminal His_6_ tag and TASOR residues 1014-1095) was acquired on a Bruker Avance 600 MHz spectrometer equipped with a triple resonance TCI cryoprobe at 298 K. Standard triple resonance experiments HNCA, CBCA(CO)NH, HNCACB, HNCO and HN(CA)CO enabled a partial resonance assignment of N- and C-terminal unstructured residues 285-291 and 368-374 of Periphilin. The largely reduced or absent signals for residues 292-367 is indicative for the increase in transverse relaxation of this structured region due complex formation with TASOR.

#### Liquid chromatography coupled to mass spectrometry (LC-MS)

Denatured Periphilin-TASOR (10 µM) was subjected to LC-MS analysis. Briefly, the complex was separated on a C4 BEH 1.7µm, 1.0 × 100 mm UPLC column (Waters, UK) using a modified NanoAcquity liquid chromatography unit (Waters, UK) to deliver a flow of approximately 50 µl min^−1.^ The column was developed over 20 min with a 2-80% (v/v) gradient of acetonitrile in 0.1% (v/v) formic acid. The analytical column outlet was directly interfaced via an electrospray ionisation source, with a hybrid quadrupole time-of-flight mass spectrometer (Xevo G2, Waters, UK). Data were acquired over a m/z range of 300–2000, in positive ion mode with a cone voltage of 30 V. Scans were summed manually and deconvoluted with MaxEnt1 (Masslynx, Waters, UK).

#### Native mass spectrometry

10 µM Periphilin-TASOR complex was buffer-exchanged into 0.1 M ammonium acetate using P6 Bio-Spin columns (BioRad). The complex was analyzed on a SYNAPT G2Si HDMS mass spectrometer (Waters, UK). Briefly, 5 µl of protein was loaded into a GlassTip emitter (New Objective, USA) and sprayed into the instrument by nano-electrospray ionization (nano-ESI) with a voltage of 1.2 kV, cone voltage 150 V, offset 150 V and trap collision energy 40 V. Scans were summed and manually deconvoluted with MassLynx4.1 (Waters, UK).

## Supplementary Figures

**Fig. EV1.**
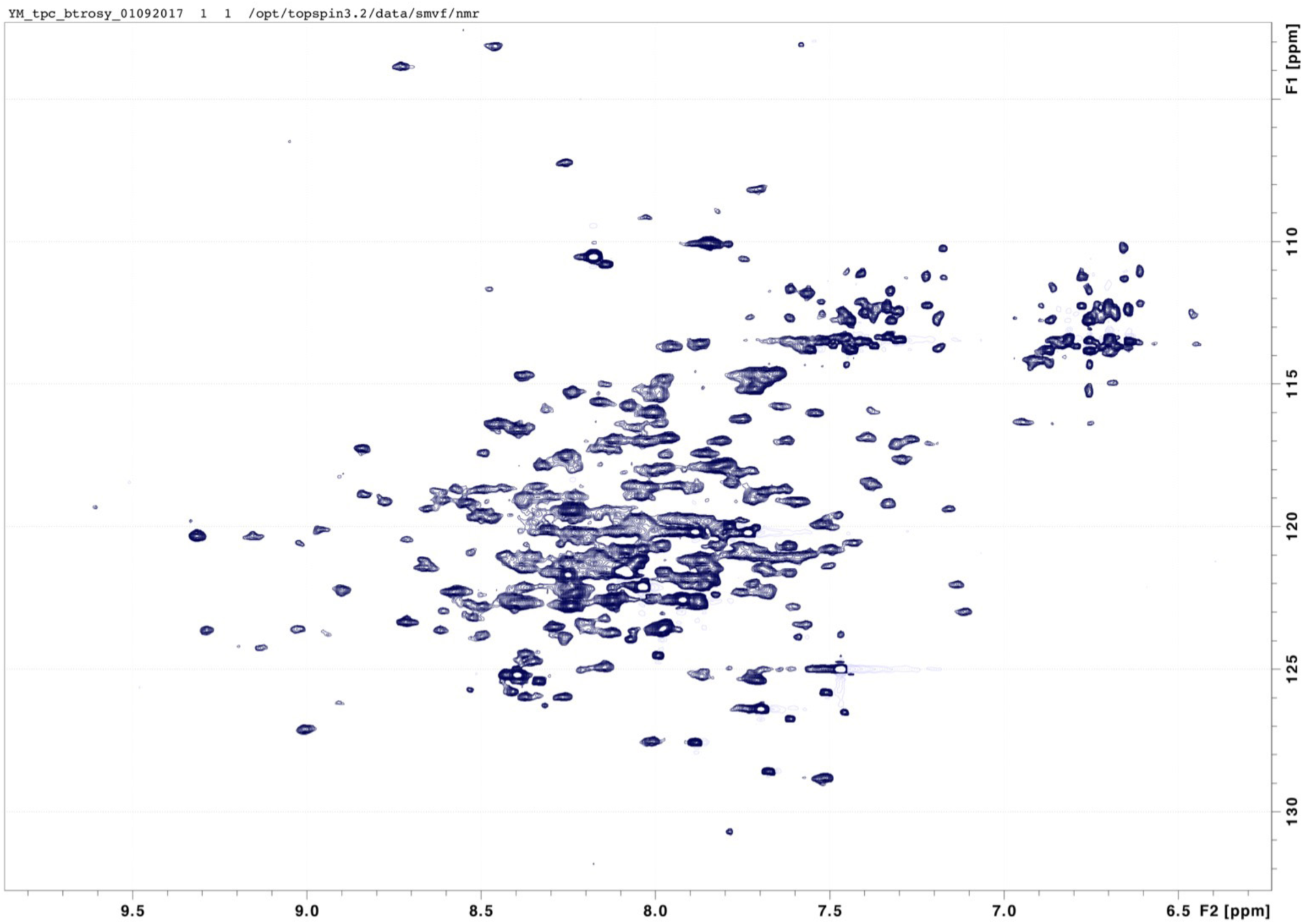
^1^H,^15^N BEST-TROSY nuclear magnetic resonance (NMR) spectrum of ^15^N,^13^C-labeled Periphilin in the Periphilin-TASOR complex. The spectrum displays backbone ^1^H,^15^N correlations for labeled Periphilin. Large intensity variations are caused by the distinctly different dynamic behavior of unstructured residues. A partial resonance assignment of intense signals in the center of the spectrum corresponding to disordered residues 285-291 and 368-374 suggested that residues 292-367 of Periphilin were ordered.

**Fig. EV2.**
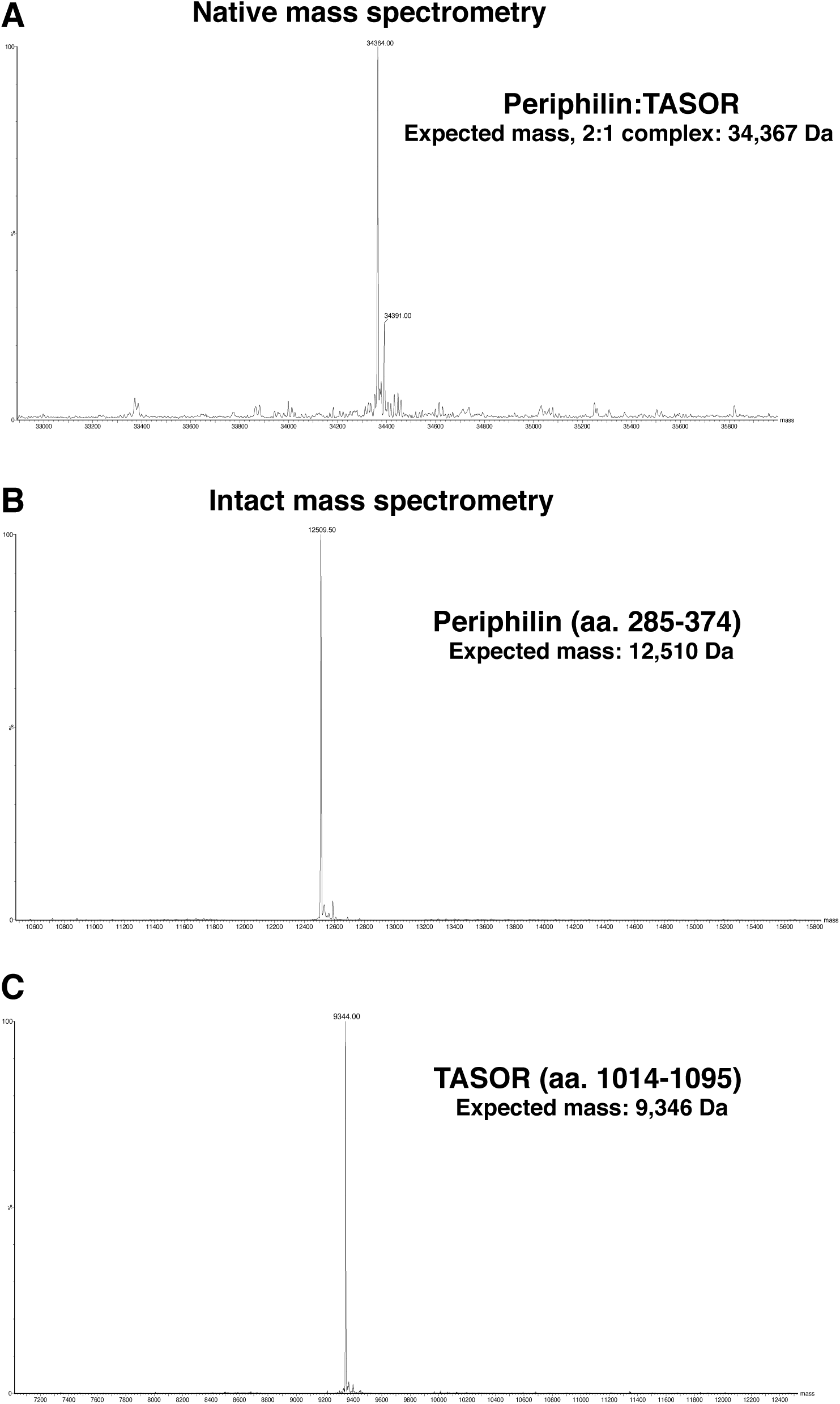
Native and intact mass spectrometry of the Periphilin-TASOR complex and its components. (**A**) Deconvoluted native mass spectrum of the Periphilin-TASOR complex collected under non-denaturing conditions. (**B**) Deconvoluted intact mass spectrum of the TASOR-binding domain of Periphilin, residues 285-374 (denaturing conditions). (**C**) Deconvoluted intact mass spectrum of the Periphilin-binding domain of TASOR, residues 1014-1095 (denaturing conditions).

**Fig. EV3.**
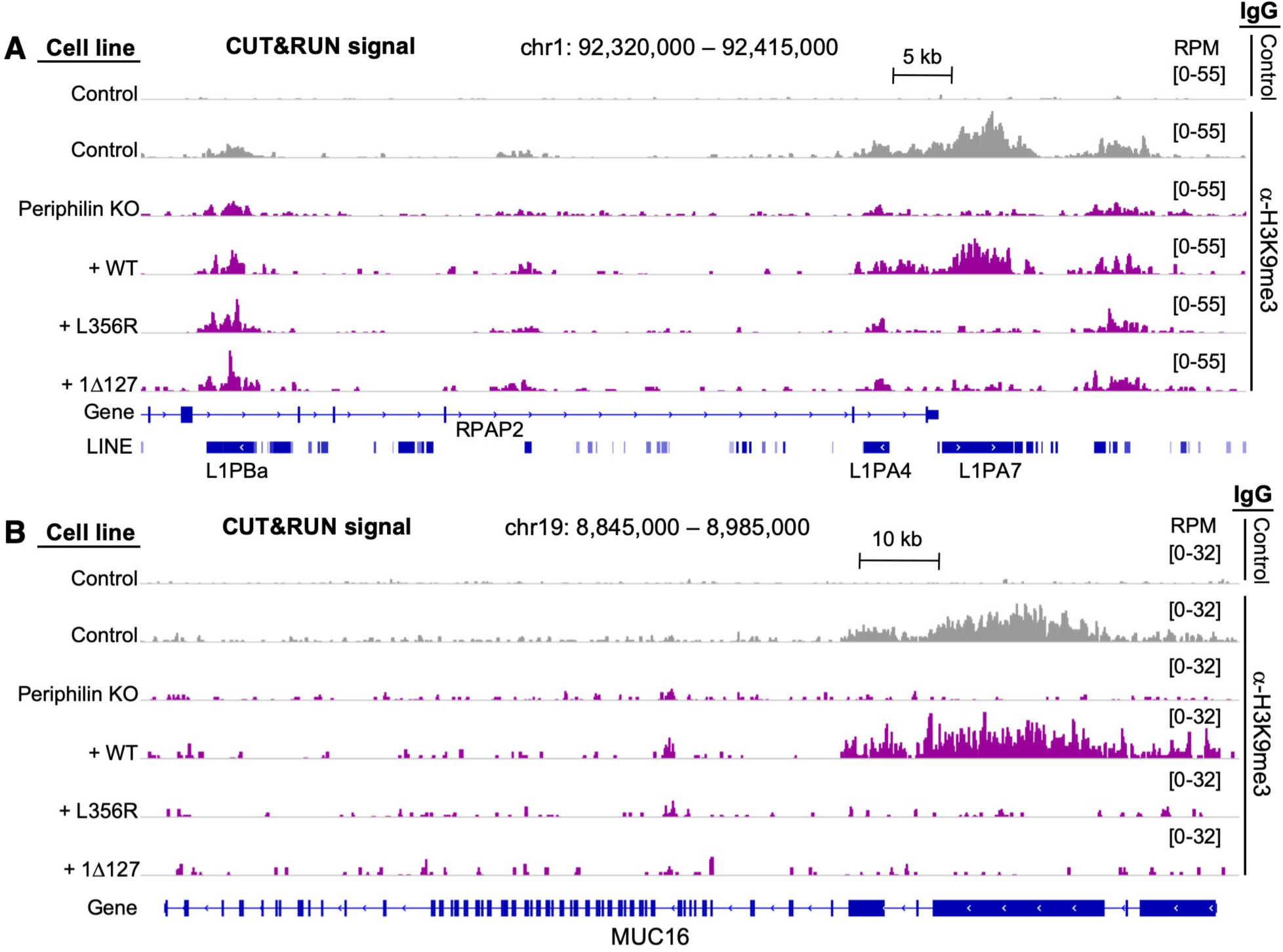
CUT&RUN genome-wide analysis of Periphilin and H3K9me3 distribution with wild-type and functionally deficient variants of Periphilin. (**A**) and (**B**) Representative snapshots of H3K9me3 distribution along the genome in the presence of different Periphilin variants. H3K9me3 distribution is shown at a full-length (6kb) L1PA7 element, (**A**), and around gene MUC16 (**B**). Both loci were shown previously to be covered in HUSH-dependent H3K9me3 (Douse *et al*, 2019). H3K9me3 from control cells and a track with a non-cognate IgG are shown in grey as positive and negative controls, respectively. The Periphilin-complemented tracks are in purple. Experiments were run in biological duplicate with similar results. RPM, reads per million, scaled to the total number of reads.

## References

Adams PD, Afonine PV, Bunkoczi G, Chen VB, Davis IW, Echols N, Headd JJ, Hung LW, Kapral GJ, Grosse-Kunstleve RW, McCoy AJ, Moriarty NW, Oeffner R, Read RJ, Richardson DC, Richardson JS, Terwilliger TC, Zwart PH (2010) PHENIX: a comprehensive Python-based system for macromolecular structure solution. Acta Crystallogr D Biol Crystallogr 66: 213–21

Alberti S, Halfmann R, King O, Kapila A, Lindquist S (2009) A systematic survey identifies prions and illuminates sequence features of prionogenic proteins. Cell 137: 146–58

Alexandrov IM, Vishnevskaya AB, Ter-Avanesyan MD, Kushnirov VV (2008) Appearance and propagation of polyglutamine-based amyloids in yeast: tyrosine residues enable polymer fragmentation. J Biol Chem 283: 15185–92

Banani SF, Lee HO, Hyman AA, Rosen MK (2017) Biomolecular condensates: organizers of cellular biochemistry. Nature reviews 18: 285–298

Castello A, Fischer B, Eichelbaum K, Horos R, Beckmann BM, Strein C, Davey NE, Humphreys DT, Preiss T, Steinmetz LM, Krijgsveld J, Hentze MW (2012) Insights into RNA biology from an atlas of mammalian mRNA-binding proteins. Cell 149: 1393–406

Chougui G, Munir-Matloob S, Matkovic R, Martin MM, Morel M, Lahouassa H, Leduc M, Ramirez BC, Etienne L, Margottin-Goguet F (2018) HIV-2/SIV viral protein X counteracts HUSH repressor complex. Nat Microbiol 3: 891–897

Chuong EB, Elde NC, Feschotte C (2016) Regulatory evolution of innate immunity through co-option of endogenous retroviruses. Science 351: 1083–7

Cooke A, Schwarzl T, Huppertz I, Kramer G, Mantas P, Alleaume AM, Huber W, Krijgsveld J, Hentze MW (2019) The RNA-Binding Protein YBX3 Controls Amino Acid Levels by Regulating SLC mRNA Abundance. Cell reports 27: 3097–3106 e5

Douse CH, Bloor S, Liu Y, Shamin M, Tchasovnikarova IA, Timms RT, Lehner PJ, Modis Y (2018) Neuropathic MORC2 mutations perturb GHKL ATPase dimerization dynamics and epigenetic silencing by multiple structural mechanisms. Nat Commun 9: 651

Douse CH, Timms RT, Tchasovnikarova IA, Protasio AV, Seczynska M, Albecka A, Prigozhin DM, Wagstaff J, Williamson JC, Dougan G, Freund S., Lehner PJ, Modis Y (2019) TASOR is a pseudo-PARP that governs assembly and co-transcriptional repression by the HUSH complex. bioRxiv

Dupressoir A, Lavialle C, Heidmann T (2012) From ancestral infectious retroviruses to bona fide cellular genes: role of the captured syncytins in placentation. Placenta 33: 663–71

Emsley P, Cowtan K (2004) Coot: model-building tools for molecular graphics. Acta Crystallogr D Biol Crystallogr 60: 2126–32

Friedli M, Trono D (2015) The developmental control of transposable elements and the evolution of higher species. Annu Rev Cell Dev Biol 31: 429–51

Friedli M, Turelli P, Kapopoulou A, Rauwel B, Castro-Diaz N, Rowe HM, Ecco G, Unzu C, Planet E, Lombardo A, Mangeat B, Wildhaber BE, Naldini L, Trono D (2014) Loss of transcriptional control over endogenous retroelements during reprogramming to pluripotency. Genome Res 24: 1251–9

Goldschmidt L, Teng PK, Riek R, Eisenberg D (2010) Identifying the amylome, proteins capable of forming amyloid-like fibrils. Proc Natl Acad Sci U S A 107: 3487–92

Goodier JL (2016) Restricting retrotransposons: a review. Mob DNA 7: 16

Greenwood EJD, Williamson JC, Sienkiewicz A, Naamati A, Matheson NJ, Lehner PJ (2019) Promiscuous Targeting of Cellular Proteins by Vpr Drives Systems-Level Proteomic Remodeling in HIV-1 Infection. Cell reports 27: 1579–1596 e7

Hancks DC, Kazazian HH, Jr. (2016) Roles for retrotransposon insertions in human disease. Mob DNA 7: 9

Holehouse AS, Das RK, Ahad JN, Richardson MO, Pappu RV (2017) CIDER: Resources to Analyze Sequence-Ensemble Relationships of Intrinsically Disordered Proteins. Biophys J 112: 16–21

Hughes MP, Sawaya MR, Boyer DR, Goldschmidt L, Rodriguez JA, Cascio D, Chong L, Gonen T, Eisenberg DS (2018) Atomic structures of low-complexity protein segments reveal kinked beta sheets that assemble networks. Science 359: 698–701

Huh JW, Kim TH, Yi JM, Park ES, Kim WY, Sin HS, Kim DS, Min DS, Kim SS, Kim CB, Hyun BH, Kang SK, Jung JS, Lee WH, Takenaka O, Kim HS (2006) Molecular evolution of the periphilin gene in relation to human endogenous retrovirus m element. J Mol Evol 62: 730–7

Hung T, Pratt GA, Sundararaman B, Townsend MJ, Chaivorapol C, Bhangale T, Graham RR, Ortmann W, Criswell LA, Yeo GW, Behrens TW (2015) The Ro60 autoantigen binds endogenous retroelements and regulates inflammatory gene expression. Science 350: 455–9

Jarvelin AI, Noerenberg M, Davis I, Castello A (2016) The new (dis)order in RNA regulation. Cell Commun Signal 14: 9

Kato M, Han TW, Xie S, Shi K, Du X, Wu LC, Mirzaei H, Goldsmith EJ, Longgood J, Pei J, Grishin NV, Frantz DE, Schneider JW, Chen S, Li L, Sawaya MR, Eisenberg D, Tycko R, McKnight SL (2012) Cell-free formation of RNA granules: low complexity sequence domains form dynamic fibers within hydrogels. Cell 149: 753–67

Kazerounian S, Aho S (2003) Characterization of periphilin, a widespread, highly insoluble nuclear protein and potential constituent of the keratinocyte cornified envelope. J Biol Chem 278: 36707–17

King OD, Gitler AD, Shorter J (2012) The tip of the iceberg: RNA-binding proteins with prion-like domains in neurodegenerative disease. Brain Res 1462: 61–80

Kurita M, Suzuki H, Kawano Y, Aiso S, Matsuoka M (2007) CR/periphilin is a transcriptional co-repressor involved in cell cycle progression. Biochem Biophys Res Commun 364: 930–6

Kurita M, Suzuki H, Masai H, Mizumoto K, Ogata E, Nishimoto I, Aiso S, Matsuoka M (2004) Overexpression of CR/periphilin downregulates Cdc7 expression and induces S-phase arrest. Biochem Biophys Res Commun 324: 554–61

Lamprecht B, Walter K, Kreher S, Kumar R, Hummel M, Lenze D, Kochert K, Bouhlel MA, Richter J, Soler E, Stadhouders R, Johrens K, Wurster KD, Callen DF, Harte MF, Giefing M, Barlow R, Stein H, Anagnostopoulos I, Janz M et al. (2010) Derepression of an endogenous long terminal repeat activates the CSF1R proto-oncogene in human lymphoma. Nat Med 16: 571–9, 1p following 579

Li W, Lee MH, Henderson L, Tyagi R, Bachani M, Steiner J, Campanac E, Hoffman DA, von Geldern G, Johnson K, Maric D, Morris HD, Lentz M, Pak K, Mammen A, Ostrow L, Rothstein J, Nath A (2015) Human endogenous retrovirus-K contributes to motor neuron disease. Sci Transl Med 7: 307ra153

Liu N, Lee CH, Swigut T, Grow E, Gu B, Bassik MC, Wysocka J (2018) Selective silencing of euchromatic L1s revealed by genome-wide screens for L1 regulators. Nature 553: 228–232

Malinovska L, Kroschwald S, Alberti S (2013) Protein disorder, prion propensities, and self-organizing macromolecular collectives. Biochim Biophys Acta 1834: 918–31

Manival X, Ghisolfi-Nieto L, Joseph G, Bouvet P, Erard M (2001) RNA-binding strategies common to cold-shock domain- and RNA recognition motif-containing proteins. Nucleic Acids Res 29: 2223–33

Matsumoto K, Meric F, Wolffe AP (1996) Translational repression dependent on the interaction of the Xenopus Y-box protein FRGY2 with mRNA. Role of the cold shock domain, tail domain, and selective RNA sequence recognition. J Biol Chem 271: 22706–12

Monahan Z, Ryan VH, Janke AM, Burke KA, Rhoads SN, Zerze GH, O’Meally R, Dignon GL, Conicella AE, Zheng W, Best RB, Cole RN, Mittal J, Shewmaker F, Fawzi NL (2017) Phosphorylation of the FUS low-complexity domain disrupts phase separation, aggregation, and toxicity. EMBO J 36: 2951–2967

Murray DT, Kato M, Lin Y, Thurber KR, Hung I, McKnight SL, Tycko R (2017) Structure of FUS Protein Fibrils and Its Relevance to Self-Assembly and Phase Separation of Low-Complexity Domains. Cell 171: 615–627 e16

Olsen JV, Blagoev B, Gnad F, Macek B, Kumar C, Mortensen P, Mann M (2006) Global, in vivo, and site-specific phosphorylation dynamics in signaling networks. Cell 127: 635–48

Olsen JV, Vermeulen M, Santamaria A, Kumar C, Miller ML, Jensen LJ, Gnad F, Cox J, Jensen TS, Nigg EA, Brunak S, Mann M (2010) Quantitative phosphoproteomics reveals widespread full phosphorylation site occupancy during mitosis. Science signaling 3: ra3

Schwartz JC, Wang X, Podell ER, Cech TR (2013) RNA seeds higher-order assembly of FUS protein. Cell reports 5: 918–25

Shin Y, Brangwynne CP (2017) Liquid phase condensation in cell physiology and disease. Science 357

Skene PJ, Henikoff S (2017) An efficient targeted nuclease strategy for high-resolution mapping of DNA binding sites. eLife 6

Skubak P, Pannu NS (2013) Automatic protein structure solution from weak X-ray data. Nat Commun 4: 2777

Soehn AS, Pham TT, Schaeferhoff K, Floss T, Weisenhorn DM, Wurst W, Bonin M, Riess O (2009) Periphilin is strongly expressed in the murine nervous system and is indispensable for murine development. Genesis 47: 697–707

Tchasovnikarova IA, Timms RT, Douse CH, Roberts RC, Dougan G, Kingston RE, Modis Y, Lehner PJ (2017) Hyperactivation of HUSH complex function by Charcot-Marie-Tooth disease mutation in MORC2. Nat Genet 49: 1035–1044

Tchasovnikarova IA, Timms RT, Matheson NJ, Wals K, Antrobus R, Gottgens B, Dougan G, Dawson MA, Lehner PJ (2015) GENE SILENCING. Epigenetic silencing by the HUSH complex mediates position-effect variegation in human cells. Science 348: 1481–1485

Timms RT, Tchasovnikarova IA, Antrobus R, Dougan G, Lehner PJ (2016) ATF7IP-Mediated Stabilization of the Histone Methyltransferase SETDB1 Is Essential for Heterochromatin Formation by the HUSH Complex. Cell reports 17: 653–659

Wang J, Choi JM, Holehouse AS, Lee HO, Zhang X, Jahnel M, Maharana S, Lemaitre R, Pozniakovsky A, Drechsel D, Poser I, Pappu RV, Alberti S, Hyman AA (2018) A Molecular Grammar Governing the Driving Forces for Phase Separation of Prion-like RNA Binding Proteins. Cell 174: 688–699 e16

Yang XJ, Zhu H, Mu SR, Wei WJ, Yuan X, Wang M, Liu Y, Hui J, Huang Y (2019) Crystal structure of a Y-box binding protein 1 (YB-1)-RNA complex reveals key features and residues interacting with RNA. J Biol Chem 294: 10998–11010

Yurkovetskiy L, Guney MH, Kim K, Goh SL, McCauley S, Dauphin A, Diehl WE, Luban J (2018) Primate immunodeficiency virus proteins Vpx and Vpr counteract transcriptional repression of proviruses by the HUSH complex. Nat Microbiol 3: 1354–1361

Zhou H, Di Palma S, Preisinger C, Peng M, Polat AN, Heck AJ, Mohammed S (2013) Toward a comprehensive characterization of a human cancer cell phosphoproteome. J Proteome Res 12: 260–71

Zhou L, Mitra R, Atkinson PW, Hickman AB, Dyda F, Craig NL (2004) Transposition of hAT elements links transposable elements and V(D)J recombination. Nature 432: 995–1001

Zhu Y, Wang GZ, Cingoz O, Goff SP (2018) NP220 mediates silencing of unintegrated retroviral DNA. Nature 564: 278–282

